# Mutual inclusivity improves decision-making by smoothing out choice’s competitive edge

**DOI:** 10.1101/2023.05.12.540529

**Authors:** Xiamin Leng, Romy Frömer, Thomas Summe, Amitai Shenhav

## Abstract

Decisions form a central bottleneck to most tasks, one that people often experience as costly. Past work proposes mitigating those costs by lowering one’s threshold for deciding. Here, we test an alternative solution, one that targets the basis for most choice costs: that choosing one option sacrifices others (mutual exclusivity). Across 5 studies (N = 462), we test whether this tension can be relieved by framing choices as inclusive (allowing selection of more than one option, as in buffets). We find that inclusivity makes choices more efficient, by selectively reducing competition between potential responses as participants accumulate information for each of their options. Inclusivity also made participants feel less conflicted, especially when they couldn’t decide which good option to keep or which bad option to get rid of. These inclusivity benefits were also distinguishable from the effects of manipulating decision threshold (increased urgency), which improved choices but not experiences thereof.

## Introduction

Humans are capable of making remarkably complex decisions, integrating over a multitude of factors and timescales^1–3^, and yet somehow even relatively banal decisions like what to order for lunch or how to word an email to a colleague can stop us in our tracks. When faced with difficult choices, we vacillate, experience persistent states of conflict and anxiety, and find ways to avoid choosing altogether, for instance by putting off choosing^4–7^ or engaging in suboptimal heuristics^8^. For many people – such as those with anxiety disorders and obsessive-compulsive disorder – these experiences of indecision and conflict can be particularly debilitating^9,10^. Whereas past work has characterized the types of decisions that are most conflicting ^6,11,12^, much less is known about how to make them less so. The primary reason for this gap is that researchers have yet to tackle the core element of choice that generates conflict in the first place: the inherent tension between selecting one option at the expense of excluding another. Here, we test whether this tension is more malleable than previously thought, and whether relieving it can improve both the outcome and the experience of decision-making.

The costs of decision-making have been extensively documented, even when selecting between ostensibly good options (“win-win choices”)^11–15^. People experience greater levels of conflict the more options they have and the more similar those options are to one another^11,16,17^. They also experience choices as more costly the higher the absolute value of those options, whether the options are all perceived to be very good or very bad^18,19^, irrespective of how similar the options or how much deliberation is required, and these costs are magnified when selecting between larger sets of high-value options ^13^. Collectively, these and other findings suggest that the source of choice costs resides in a simple fact that permeates all of decision making: that when choosing one option we have to sacrifice all others, that is, that our choices are *mutually exclusive* of one another (e.g., we must ultimately settle on a subset of our options for lunch, sending an email, and so on). This mutual exclusivity creates a tension whereby a person feels a tug towards and against each of their options (what Miller^12^ referred to as ‘double approach-avoidance’ because acquiring one outcome means losing out on another), and this tension intensifies the more valuable the potential gains (and conversely the potential losses).

A prominent approach to resolving this conflict has revolved around how a person sets their threshold for deciding, which defines the weight they place on speed (decision time) versus accuracy (choosing the best option in a set). For instance, rather than trying to select the best possible option, a person can choose the first option that meets a certain set of criteria (satisficing)^20^, an approach that has been shown under certain conditions to correlate with improved psychological wellbeing^21,22^ (but see^23,24^). A related solution involves allowing one’s decision threshold to decrease (collapse) over the course of a decision, setting progressively lower standards for identifying an option as the “best” until ultimately one is effectively chosen at random^25^. Indeed, previous work has shown that decision-makers can become more productive (i.e., make more decisions per unit time) when tighter choice deadlines are enforced, forcing them to dynamically decrease their threshold to meet a given deadline^26^.

Threshold adjustments provide a sensible resolution to difficult decisions because they can be controlled explicitly by the decision-maker (and/or socially engineered by their environment through deadlines) and they can guarantee that a choice is ultimately made without substantial opportunity cost of time^27,28^. However, lower thresholds also necessarily come with the cost of a potential sacrifice to choice accuracy^29^. Moreover, in part because of these potential declines in accuracy and their potential for inducing feelings of urgency and post-choice regret, such threshold adjustments may have limited benefit (and potential added detriment) for the subjective experience of choosing^30,31^. Threshold adjustments thus offer a stopgap for limiting the costs of decision-making, but they fail to address the push-pull relationship between choices that is believed to give rise to these costs, in part because they are offered under the assumption that this competition reflects an immutable property of choice. What if this property is not in fact so immutable?

Here, across 4 studies, we test the possibility that a person’s perception of the competition between their options can be altered in such a way that the person can weigh their options more independently, and that this can result not only in experiences of less choice conflict but also in all-around better choices. To do so, we have participants choose their favorite option out of a choice set, under conditions where the other options will no longer be available after (*exclusive* choice) and under conditions where they can go on to select other options from that set (*inclusive* choice). Despite this added flexibility, we show that participants still choose their favorite option first in the inclusive condition, and they do so *more efficiently* than in the exclusive condition. We show that these and other patterns of choice behavior from our experiment are selectively accounted for by a computational model in which choice inclusivity reduces the level of competition (mutual inhibition) between potential responses. We further show that manipulations of choice inclusivity generate distinct behavioral patterns from, and confer unique benefits relative to, changes in choice urgency resulting from tighter deadlines. Most notably, unlike urgent choices, inclusive choices feel *less conflicting* than their alternative (i.e., non-urgent exclusive choice). Tying this work to recent studies identifying the conditions under which choice costs are greatest^13,19,32^, we show that this beneficial impact of inclusivity on the experience of choosing varies as a function of (a) the overall value of the choice set and (b) whether choosing which options to acquire versus which to remove. Collectively, our findings provide a comprehensive account of how and why decision-making can be improved by increasing the inclusivity of one’s choices.

## Results

To test the influence of choice exclusivity on decision making, we sought to relax the constraint that people can only choose one option from a given choice set, and to compare between choices with and without this constraint. To achieve this goal, we designed a value-based decision making task involving a series of choices between sets of four consumer products (Figure 1). On each trial, participants in Study 1 (N=82; see Methods and Materials) were asked to select their favorite of these four options. For half of these trials (*exclusive* choices), this was the only choice participants made from the set; for the other half of trials (*inclusive* choices), participants were allowed to subsequently choose as many additional options from the set as they preferred. Exclusive and inclusive choices were interleaved throughout the session and were explicitly cued by a colored fixation cross prior to and throughout the trial. Critically, irrespective of the choice type, participants were always told to select the best item first. Choice sets were constructed to vary in the overall (mean) value (OV) and relative value (quantified as the difference between the value of the highest-rated product and the mean value of the remaining products; RV) of the options, based on item-wise ratings given by participants earlier in the session (see Methods). After making all of the choices, participants viewed each choice set again and retrospectively rated the level of choice conflict they had experienced while engaging in that choice ^13,19^.

**Figure 1.**
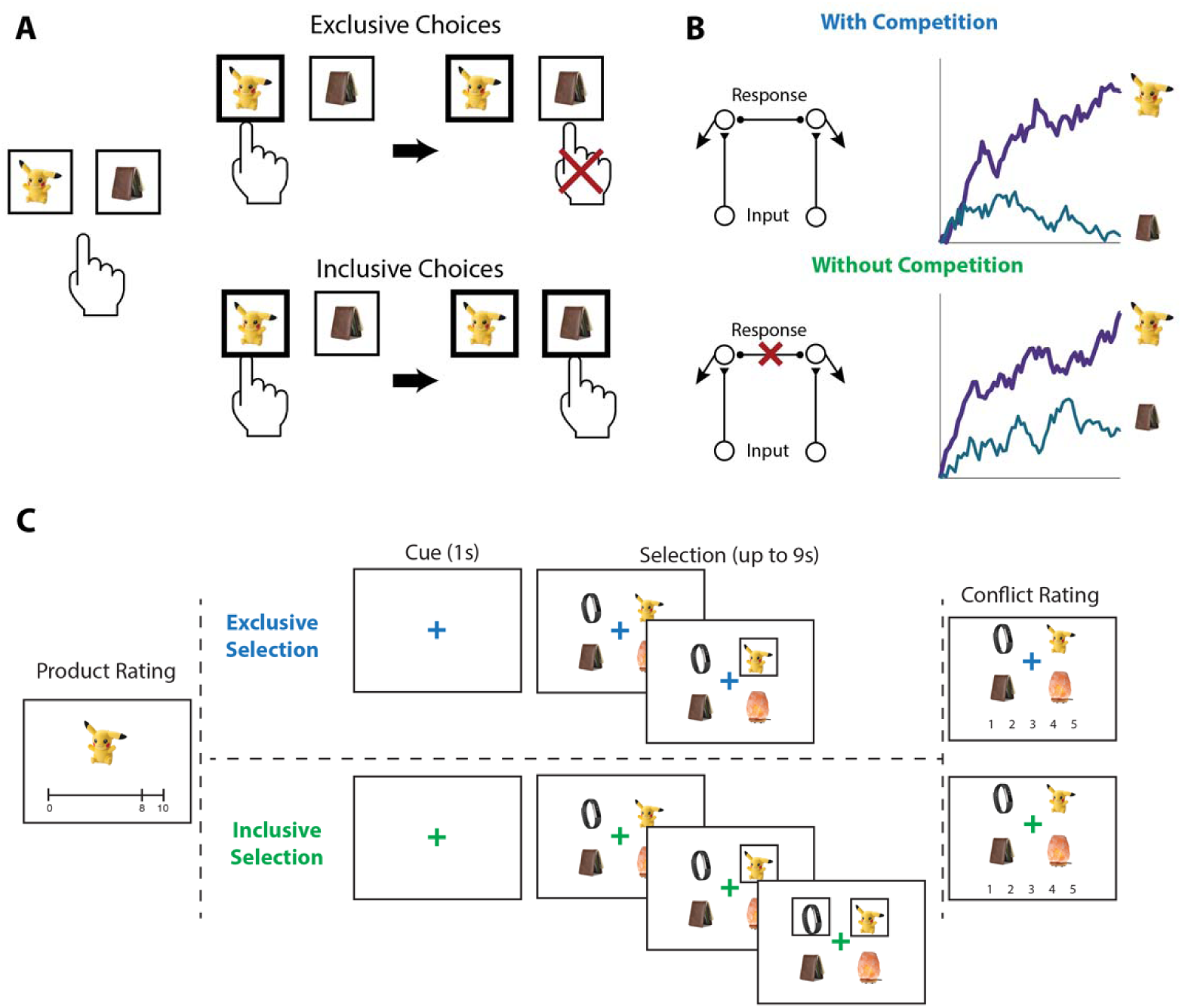
Illustration of choice inclusivity, model demonstration, and task paradigm. **(A)** Illustration of choice inclusivity. When choices are exclusive, choosing one option excludes the opportunity of choosing the others. Allowing choosing the other options in subsequent choices induces choice inclusivity. **(B)** Demonstration of the Leaky Competitive Accumulator model (LCA). With competition between options (top; as in the exclusive choices), evidence for the winning alternative will ramp up and suppress accumulation of evidence for the remaining options. Without competition (bottom; as in the inclusive choices), evidence for all alternatives will ramp up independently. **(C)** Task paradigm. Participants individually rated a series of products on how much they would like to have each one and subsequently saw sets of four products and were asked to choose the one they like best. On exclusive choice trials, the trial then ended. On inclusive choice trials, participants were allowed to select as many additional products as they liked. Finally, participants saw all option sets again and rated the level of conflict they experienced when making their choices.

### Inclusive choices are more efficient

Consistent with previous studies, we found that exclusive choices were faster and more accurate the greater the difference between the best option and the average value of the remaining options (i.e., with higher *relative value*; reaction time RT: β*_RV_*=-0.19, 95% CI=[-0.23, -0.15], p<0.001, Figure 2C; Accuracy: log-odd*_RV_*=0.68, 95% CI=[0.59, 0.78], p<0.001, Figure 2D). Comparing these choices to the first choice in the inclusive condition, we found that inclusive choices were significantly faster (M*_excl_*= 2.87s; M*_incl_* =2.57s; β*_incl_*=-0.30, 95% CI=[-0.38,-0.22], p<0.001; Figure 2A). While this might at first suggest that participants were simply making this initial choice at random when it was inclusive – with the understanding that they could subsequently choose as many additional items as they wanted from the set – this was not in fact the case. We found that the likelihood of choosing the best item first (choice “accuracy”) decreased more modestly between exclusive and inclusive choices (M*_excl_*=0.49; M*_incl_* = 0.47; log-odd*_incl_*=-0.10, 95% CI=[-0.19,-0.01], p=0.029; Chance level of accuracy: 0.25; Figure 2A), and that accuracy was equally sensitive to the relative value of one’s options in both conditions (relative value by inclusivity: log-odd*_RV_*_×*incl*_=-0.06, 95% CI=[-0.17, 0.04], p=0.246, Figure 2D). These findings suggest that, despite being faster, participants were discriminating between the values of their options similarly well when making inclusive relative to exclusive choices.

**Figure 2.**
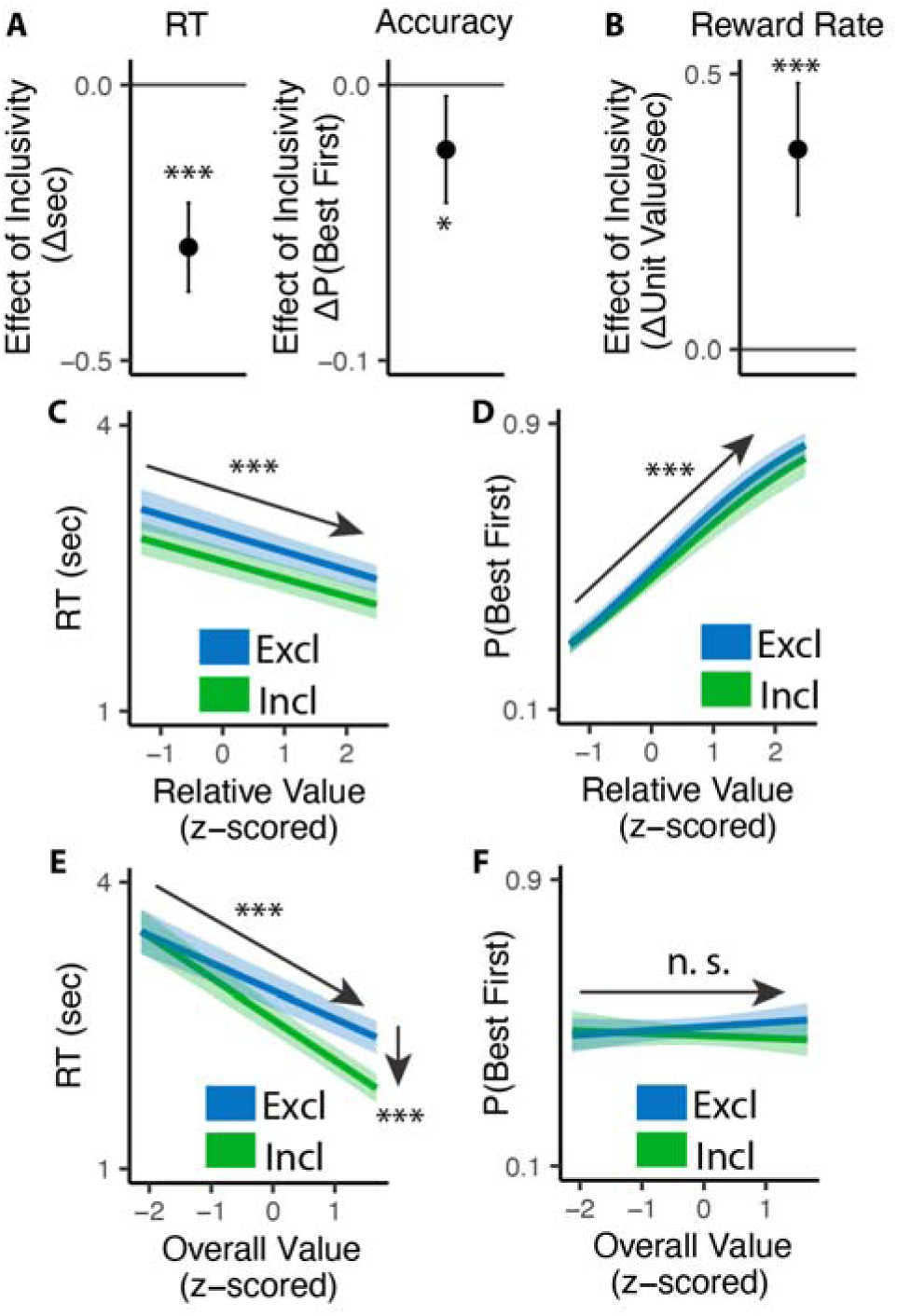
Influence of choice inclusivity on speed and accuracy during initial choices. **(A-B)** Compared to exclusive choices, people made faster and slightly less accurate (A) decisions in inclusive choices, achieving higher reward rate (B). **(C-D)** People choose faster and more accurately the greater the difference between the best option and the others. **(E-F)** People were faster to choose when the overall value of a choice set was higher. This speeding effect was greater for inclusive relative to exclusive choices. Overall value did not significantly influence choice accuracy. These effects did not differ across conditions. Error bars and shaded areas indicate 95% confidence intervals. n.s.: p>0.05; *: p<0.05; ***: p < 0.001.

Collectively, these patterns suggest that participants were overall more efficient in making inclusive relative to exclusive choices: choosing quickly but effectively. To quantify this change in efficiency, we calculated the reward rate accrued (hypothetically) for each condition by dividing the value of the chosen item by the time taken to make a given response, confining to the initial choice on each trial. We found that participants achieved a significantly higher reward rate for these initial choices when choices were inclusive (M*_exl_*=3.12, M*_incl_*=3.48; β*_incl_*=0.36, 95% CI=[0.24,0.48], p<0.001; Figure 2B). In other words, each unit time spent choosing was more productive when choice exclusivity was relaxed.

A final key difference emerged between choice behavior in these conditions, which provided important clues as to underlying computational mechanisms. As in previous work, when participants were making exclusive choices their response times were negatively correlated with the overall (average) value of the choice set (i.e., they were faster when their options were overall more valuable: β*_OV_*=-0.29, 95% CI: [-0.34, -0.24], p<0.001; Figure 2E). When participants were making inclusive choices, by contrast, this negative slope became much steeper, exacerbating the speeding effect of overall value on choice RTs (β*_OV_*_×*incl*_=-0.14, 95% CI: [-0.20, - 0.08], p<0.001; Figure 2E). These RT effects were not mirrored in accuracy - overall value did not influence choice accuracy overall (exclusive choices: log-odd*_OV_*=0.05; 95% CI: [-0.03, 0.13]; p=0.243; Figure 2F), nor did it interact with choice inclusivity (log-odd*_OV_*_×*incl*_=-0.08; 95% CI: [-0.19, 0.03]; p=0.148; Figure 2F). The behavioral patterns in Study 1 are replicated in a follow-up study with temporally separated initial and subsequent choices in the inclusive condition (Figure S1 in Supplementary Materials) and a subset in this study with incentive-compatible settings (Figure S2 in Supplementary Materials).

### The benefits of choice inclusivity are uniquely accounted for by reductions in mutual inhibition

We predicted that these inclusivity-related changes in choice behavior would be accounted for by differences in competition between options, which was instantiated in our model as the level of mutual inhibition between potential responses. However, a plausible alternative to this – which has been the focus of past work on choice simplification^26^ – is that participants were instead lowering their response threshold when faced with inclusive choices relative to exclusive choices. Such strategic threshold-lowering has been demonstrated empirically in other research^33^, and can take either of two forms: an overall decrease in one’s threshold for responding, and/or a sharper decrease (collapse) in an initial threshold over the course of a choice (Figure 3B). To adjudicate between these different mechanistic accounts, we compared our empirical findings to patterns of choice behavior predicted by simulations of a Leaky Competing Accumulator model (LCA)^34,35^ when varying (a) mutual inhibition, (b) the height of an initial response threshold, and (c) the rate at which that threshold collapses (see Methods and Materials).

**Figure 3.**
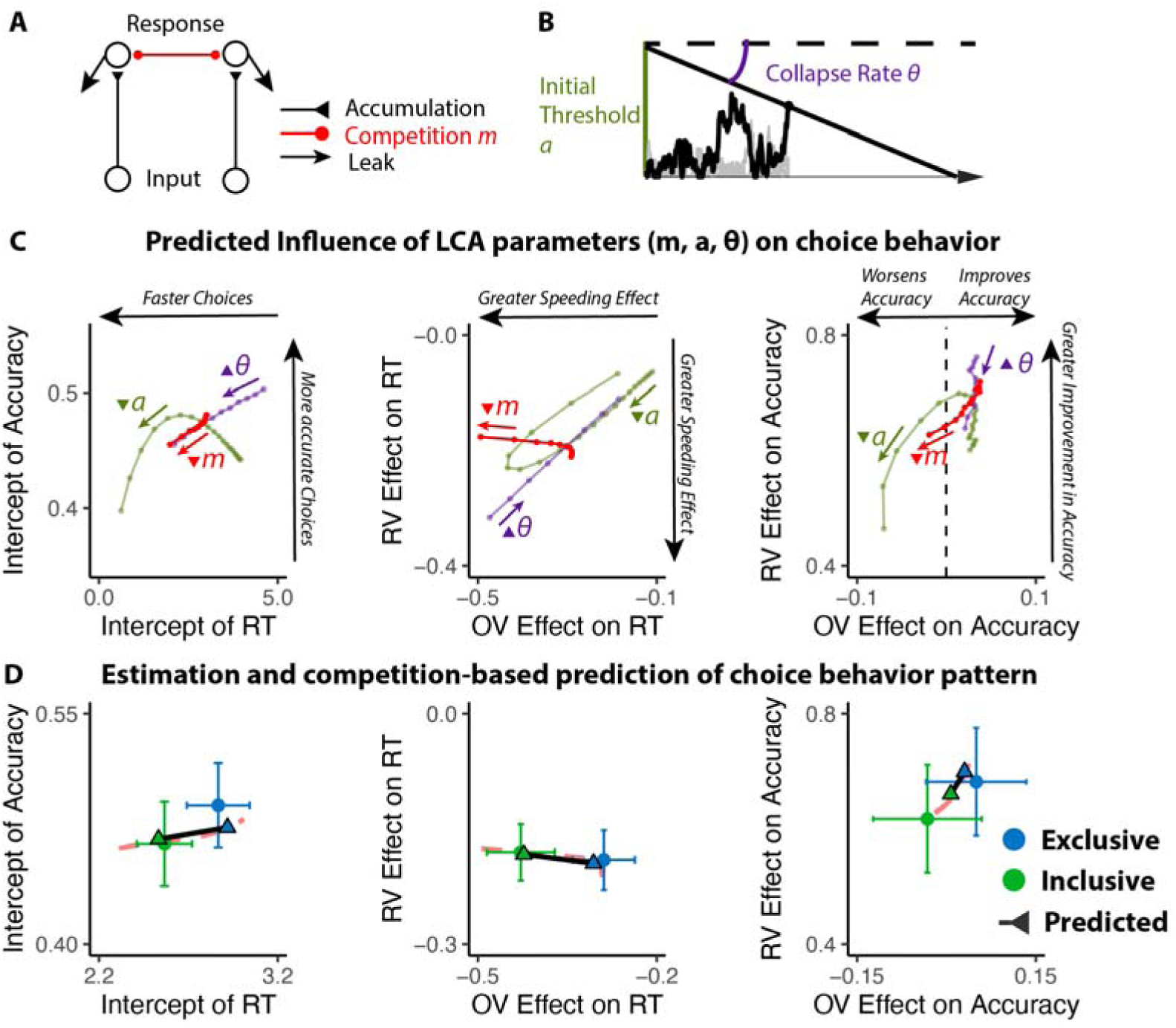
Influences of mutual inhibition based on LCA simulations. **(A-B)** Schematic and a sample iteration of the collapsing-boundary LCA. **(C)** Decrease in mutual inhibition *m* but not decrease in initial threshold *a* and/or increase in collapse rate *ϑ* predicts the observed influence of choice inclusivity on overall RT and accuracy, and the effects of relative value (RV) and overall value (OV). Each dot reflects regression coefficients/intercepts based on the average of 100 iterations. The upward triangles indicate increase in magnitude, whereas the upside-down triangle indicates decrease in magnitude. **(D)** Simulations (black lines) capture the empirical patterns observed in both conditions, dashed red lines represent predicted influence of competition in panel C.

Our simulations revealed that qualitatively different patterns of choice behavior should emerge when varying mutual inhibition versus response threshold (Figure 3C). First, whereas both forms of adjustment should enhance speeding effects of overall value, reductions in threshold and increase in collapse rate should produce correlated enhancements in value-difference related speeding but reductions in mutual inhibition should not strongly affect the relationship between relative value and RT. Second, reductions in threshold and increase in collapse rate should strongly reduce value-difference related change in accuracy, but reductions in mutual inhibition should not produce such a strong effect. In each of these cases, our empirical data was consistent only with mutual-inhibition-related predictions and not the threshold-related predictions (Figure 3D; also see Figure S3 and Table S1 in Supplementary Materials).

### Inclusive choices feel less conflicting

Our findings show that people make choices more efficiently when framed in an inclusive rather than exclusive choice setting. To test whether differences in choice inclusivity can further alter a person’s *experience* of choosing, at the end of the experiment we had participants retrospectively rate the level of choice conflict they experienced while making each of the choices^19^. We found that participants experienced less choice conflict when making inclusive choices than when making exclusive ones (M*_excl_* = 2.61; M*_incl_* = 2.15; β*_incl_*=-0.46, 95% CI=[-0.62, - 0.29], p<0.001; Figure 4A).

**Figure 4.**
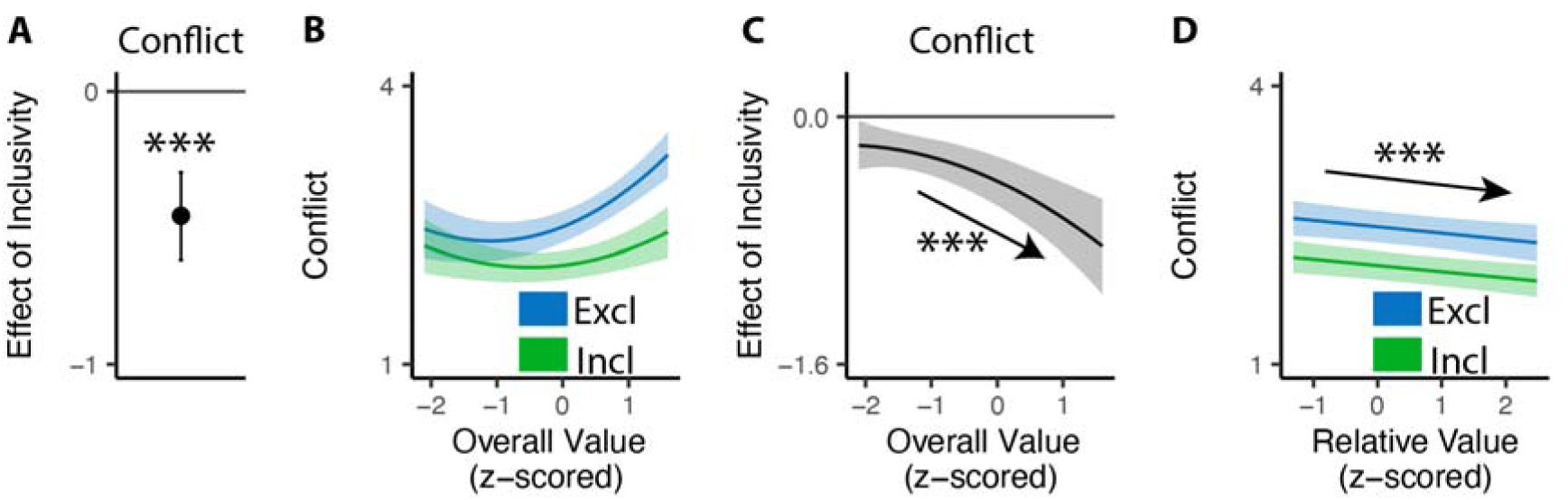
Influences of choice inclusivity on the subject experience of conflict. **(A)** People experience a higher level of conflict in exclusive choices compared to inclusive ones. **(B)** There is a typical U-shaped relationship between the overall value of a choice set and level of conflict in exclusive choices (blue) but the level of conflict is reduced in inclusive ones especially when the overall value is higher (green). **(C)** The difference between exclusive and inclusive choices on conflict increases with overall value. **(D)** The difference between exclusive and inclusive choices does not vary with relative value. The shaded areas indicate 95% confidence intervals. ***: p<0.001.

Notably, this reduction in choice conflict for inclusive choices was not uniform across choices, but rather varied with the overall value of one’s options. Consistent with previous studies ^13,19,36^, we found that choice conflict exhibited a U-shaped relationship with overall value when controlling for relative value (also see Figure S4 in Supplementary Materials): greatest when choosing among options that are especially high in value (inducing high levels of conflict over which was *most preferred*) or especially low in value (inducing high levels of conflict over which option was *least unpreferred*) (exclusive choices: β*_OV_linear_*=24.11, 95% CI=[12.95, 35.27], p<0.001; β *_OV_quad_*=15.07, 95% CI=[9.54, 20.61], p<0.001; Figure 4B). We found that the shape of this curve changed when participants were making inclusive choices. Specifically, the decreases in choice conflict we report above (when collapsing across all trials) were greatest when participants were choosing among higher value options and smallest when participants were choosing among lower value options (β*_incl_*_×*OV_linear*_=-18.76, 95% CI=[-27.07,-10.45], p<0.001; β*_incl_*_×*OV_quad*_=-4.87, 95% CI=[-9.97, 0.23], p=0.061; Figure 4C; also see Table S8 and Figure S4 in Supplementary Materials). Thus, the benefit of inclusivity on experiences of choice conflict increases with overall value. By contrast, no such interaction was found between choice inclusivity and the influence of *relative* value on choice conflict (β*_incl_*_×*RV*_=-0.00, 95% CI=[-0.05,0.05],p=0.894; Figure 4D).

### The benefits of choice inclusivity extend to the removal of bad options

In Study 1, we found that when people knew they would be able to select additional options from a set (inclusive choices), they felt less conflicted and chose more efficiently. Interestingly, choice inclusivity led to reduced choice conflict for most choices, but not when choosing between the least valuable options. This pattern is consistent with previous work suggesting that conflict for low-value choices stems from not wanting any of the options^19^ as participants were required to choose at least one of these options for both conditions. By contrast, high-value choice conflict – which stems from desire to select more than one option, could be alleviated by enabling participants to choose as many options as they want.

To test this account, and to rule out other contributions to these behavioral findings related to the salience of the rewards themselves^32,37–39^, Study 2 (N=98; see Methods and Materials) inverted the choice framing in Study 1. Rather than choosing which option(s) they wanted to *select*, we instead had participants assume a set of options had already been selected for them and asked them to choose which option(s) they wanted to *de-select* (i.e., remove) from that set (Figure 5A). Analogous to Study 1, they were always asked to first choose the item they most wanted to remove, and in inclusive choices were subsequently allowed to remove as many of the other options as they wanted. We predicted that choice inclusivity would impact behavior the same way as in Study 1, but that it would result in greatest reduction in choice conflict for the choice set with least valuable options rather than the most valuable ones.

**Figure 5.**
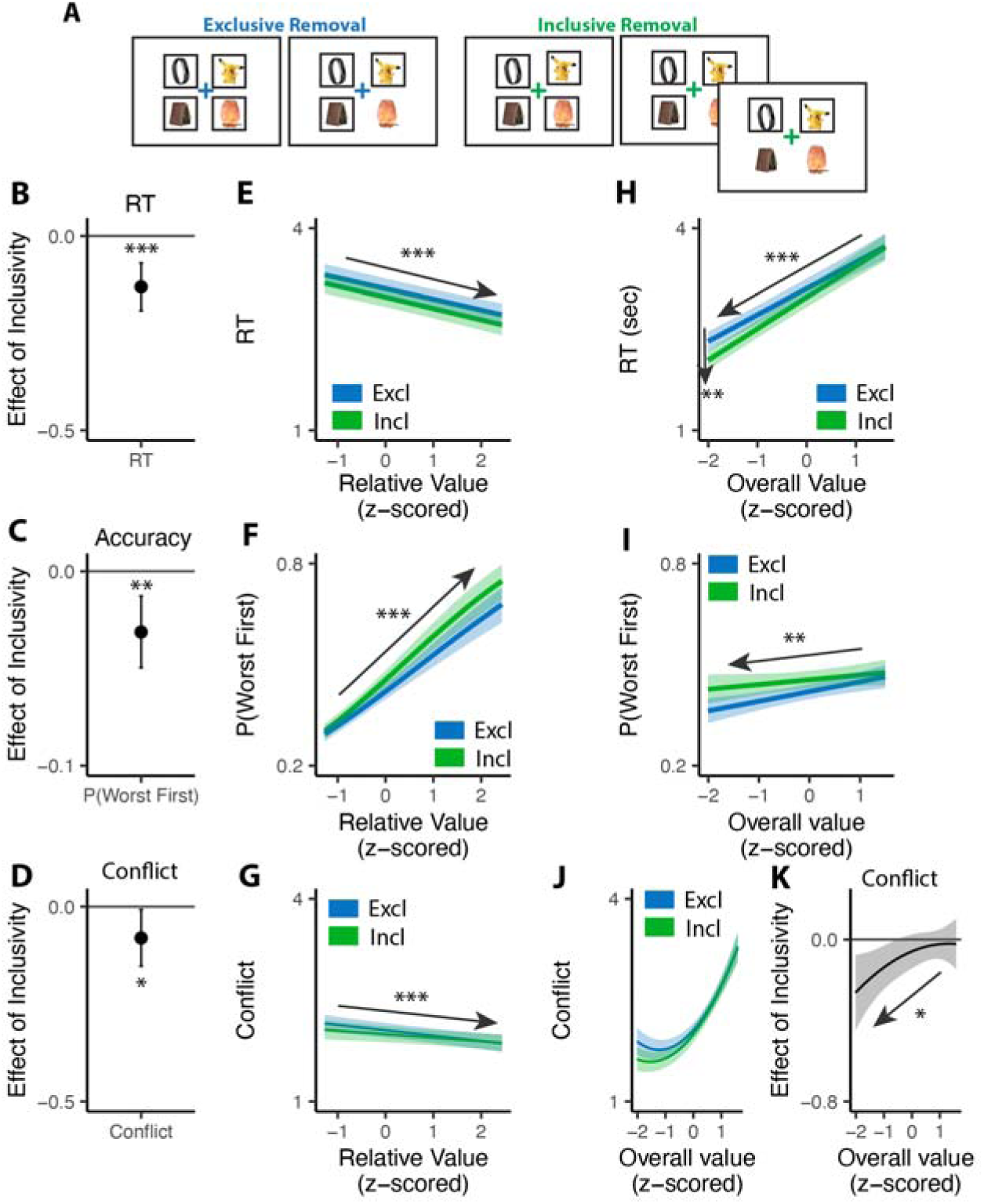
Influence of choice inclusivity when removing options. **(A)** In the removal task, participants saw sets of four products that were pre-selected (indicated by black frames) and were asked to remove the one they liked least. On exclusive choice trials, the trial then ended. In inclusive choice trials, participants were allowed to remove as many additional products as they liked. The product rating and conflict rating phases followed the same settings in Study 1. **(B-C)** Compared to exclusive removals, people made faster and similarly accurate decisions in inclusive ones. **(E-F)** People remove faster and more accurately the greater relative value (e.g., the difference between the worst option and the others). The effect on RT did not differ across conditions but the effect on accuracy decreases in inclusive cases. **(H-I)** People were faster but less accurate to remove when the overall value of a choice set was lower. The speeding effect was greater for inclusive relative to exclusive removals. The effect of overall value on accuracy is similar between two kinds of removals. **(D)** People experience a higher level of conflict in exclusive removals compared to inclusive ones. **(G)** The level of conflict decreases with relative value and the effect of relative value does not differ between conditions. **(J)** The U-shaped relationship between the overall value of a choice set and level of conflict is reduced in inclusive compared to exclusive removals, whereby conflict is specifically reduced for low-value choice sets. **(K)** The difference between exclusive and inclusive choices on conflict decreases with overall value. Error bars and shaded areas indicate 95% confidence intervals. *: p<0.05; ***: p<0.001.

Consistent with past research, when the choice goal is to remove the least preferred option, the effect of overall value on speed is reversed - participants are faster when the overall value is lower (RT in exclusive: β*_OV_*=0.39, 95% CI=[0.35,0.44], p<0.001). Both of these predictions were confirmed. First, just as participants choosing which option they most wanted from a set (Study 1), participants choosing which item they most wanted to remove from a set were faster (M*_excl_*=3.11, M*_incl_*=2.97; β*_incl_*=-0.13, 95% CI=[-0.19,-0.07], p<0.001; Figure 5B) and exhibited a stronger effect of overall value on choice speed (β*_incl_*_×*OV*_=0.07, 95% CI=[0.02,0.12], p=0.003; Figure 5H) under an inclusive framing. These and other patterns of choice behavior (e.g., no interactions between exclusivity and relative value effects) were again uniquely accounted for by an accumulator model with varying levels of mutual inhibition (Figure 5E).

Second, as in Study 1, we found that participants experienced less conflict overall when engaged in inclusive relative to exclusive choices (M*_excl_*=2.25, M*_incl_*=2.17; β*_incl_*=-0.08, 95% CI=[-0.15,-0.01], p=0.030; Figure 5D), and once again found that this effect varied based on the overall value of the choice set. More importantly, as predicted, this relationship was opposite to the one we found in Study 1 (i.e., negative rather than positive; β*_incl_*_×*OV_linear*_=7.23, 95% CI=[1.23, 13.24], p=0.018; β*_incl_*_×*OV_quad*_=-2.45, 95% CI=[-7.29,2.39], p=0.321; Figure 5J-K). When participants were deciding between *high-value* options, choice inclusivity diminished their experience of choice conflict when choosing which one to *select* (Study 1) but not when choosing which one to *remove* (Study 2). Conversely, when deciding between *low-value* options, relaxing choice exclusivity diminished their experience of choice conflict when choosing which one to *remove* (Study 2) but not when choosing which one to *select* (Study 1).

### The benefits of choice inclusivity persist in the absence of time pressure

Studies 1-2 validate our core predictions regarding the effect of choice inclusivity on behavior and subjective experience, under conditions where participants select options to obtain (Study 1) and de-select options that they would like to remove (Study 2). One potential concern, though, is that participants in both studies were given a limited time window to respond (9 s) which, while reasonably long for a single choice, may have introduced additional time pressure when participants were able to select up to four options (inclusive choices). Previous work suggests that such time pressure can result in a dynamically decreasing response threshold ^40^ which could have contributed to the differential patterns of behavior and conflict we observed across the two conditions. To rule this out, Studies 3A and 3B (Ns = 59 and 61; see Methods and Materials) replicated the procedures in Studies 1 and 2, respectively, but omitted the choice deadline. These studies also included more up-to-date products, but were otherwise identical to the studies above. Our results confirm our previous behavior findings in Study 1 and 2 (Figure 6A; see Figure S5-6 and Table S9-11 in Supplementary Materials). We also observed that the influence of inclusivity depends on both the choice action (selection vs. removal) and the overall value of choice set (Figure 6B; see Figure S7-8 and Table S12 in Supplementary Materials) in the same direction as we see in Study 1 and 2.

**Figure 6.**
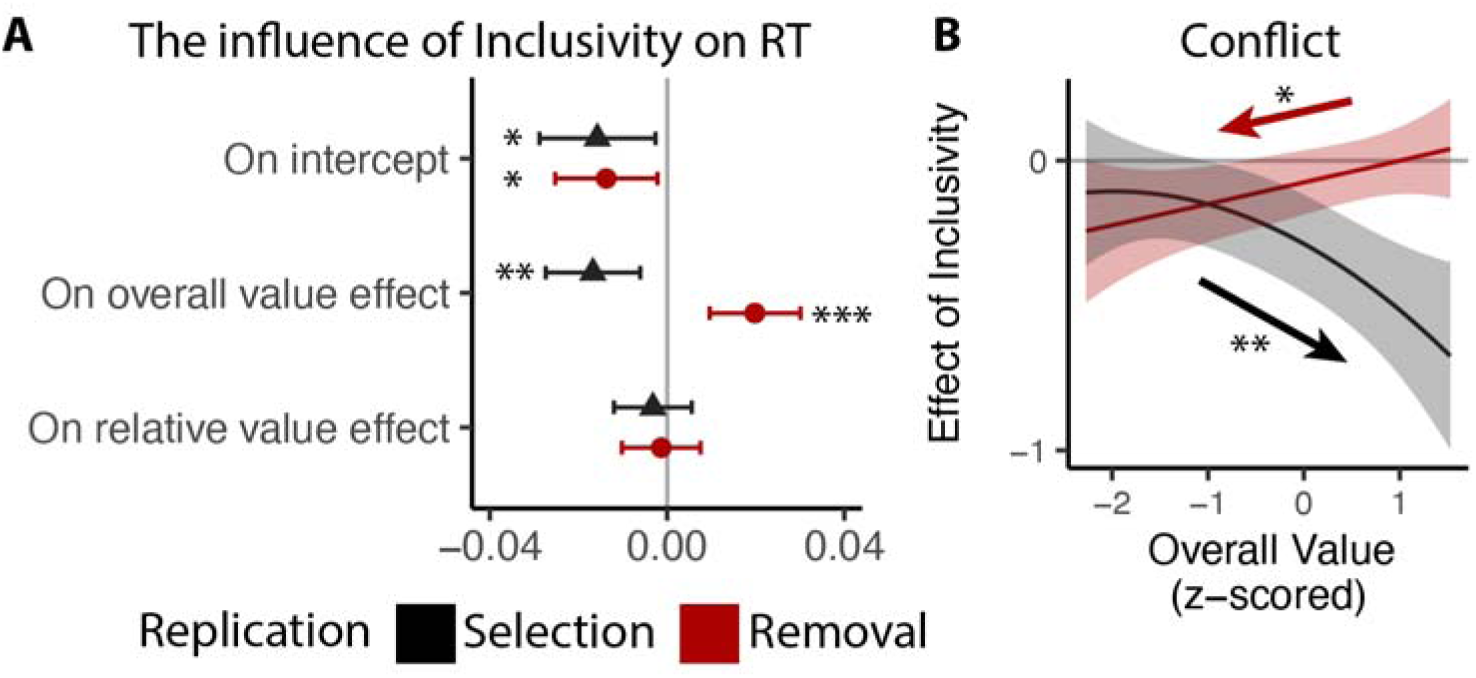
Replication of choice inclusivity effects on choice behavior during selection and removal. We followed the same analysis procedure as in Study 1 and 2, except that to account for the long tail in RTs without the time limit, we log-transformed RTs prior to the analysis. Across replication studies for selection and removal, we confirmed the findings that: **(A)** participants were faster with stronger effect of overall value on the reaction time in inclusive choices; and **(B)** the effect of inclusivity on the conflict increases with high/low overall value in selection/removal task. Error bars and shaded areas indicate 95% confidence intervals. *: p<0.05; **: p<0.01; ***: p<0.001.

### Choice inclusivity confers unique benefits relative to choice urgency

Previous work suggests that decisions can be optimized by tightening one’s decision deadline, constraining their natural inclination towards setting their decision thresholds too high (relative to what would be reward rate-optimal)^26,33^. As we show in Studies 1-3, our manipulation of choice exclusivity optimizes decision-making by altering a different decision parameter (mutual inhibition), thus generating qualitatively different patterns of choice behavior than what would be expected from threshold adjustments (Figure 3), and which persist in the absence of time pressure (Figure 6). While these findings establish that inclusivity serves as an alternate path to optimizing choice relative to changes in choice threshold/urgency, it is unclear whether these paths reach similar or different endpoints (i.e., qualitatively similar improvements in decision-making). To test the extent to which these two forms of choice optimization yield comparable effects, we had a separate group of participants (Study 4, N=85; see Methods and Materials) perform the same experiment as in Study 1 but rather than varying choice exclusivity we instead had them always make exclusive choices and instead varied whether this was done under *high urgency* (3s choice deadline) or *low urgency* (no time limit, comparable to exclusive choices in Study 1).

Consistent with our model simulations (Figure 3C), urgency (which we predicted would lead to reductions in decision threshold) produced qualitatively distinct changes in choice behavior than inclusivity (which we predicted would lead to reductions in mutual inhibition). When having to respond under higher choice urgency, participants were both faster (M*_low urgency_*=2.66, M*_high urgency_*=1.73; β*_urgency_*=-0.93, 95% CI=[-1.10,-0.75], p<0.001) and less accurate (M*_low urgency_*=0.46, M*_high urgency_*=0.43; log-odd*_urgency_*=-0.14, 95% CI=[-0.23,-0.05], p=0.003). Consistent with previous demonstrations of urgency’s utility for choice optimization ^26^, these changes collectively led to an overall higher reward rate on high urgency trials (M*_high urgency_*=4.44) relative to low urgency ones (M*_low urgency_*=3.55) (β*_urgency_*=0.88, 95% CI=[0.75,1.02], p<0.001). These changes in overall choice performance are directionally similar to those observed when varying choice inclusivity, but our simulations predict a key dissociation when examining the influence of choice value on behavior (Figure 3C): whereas changes in mutual inhibition should selectively enhance the speeding effect of overall value on RT, changes in threshold should *diminish* this speeding effect similarly for both overall *and* relative value. Both of these predictions were confirmed: choice urgency reduced the speeding effects of overall value and relative value with similar magnitude (β*_urgency_*_×*OV*_=0.23, 95% CI = [0.16, 0.30], p<0.001; β*_urgency_*_×*RV*_=0.23, 95% CI = [0.16, 0.30], p<0.001; Figure 7A-B; see also Figure S9 in Supplementary Materials). These findings establish that inclusivity versus urgency exert dissociable influences on mutual inhibition versus decision threshold, and demonstrate the utility of each as a potential choice optimization tool.

**Figure 7:**
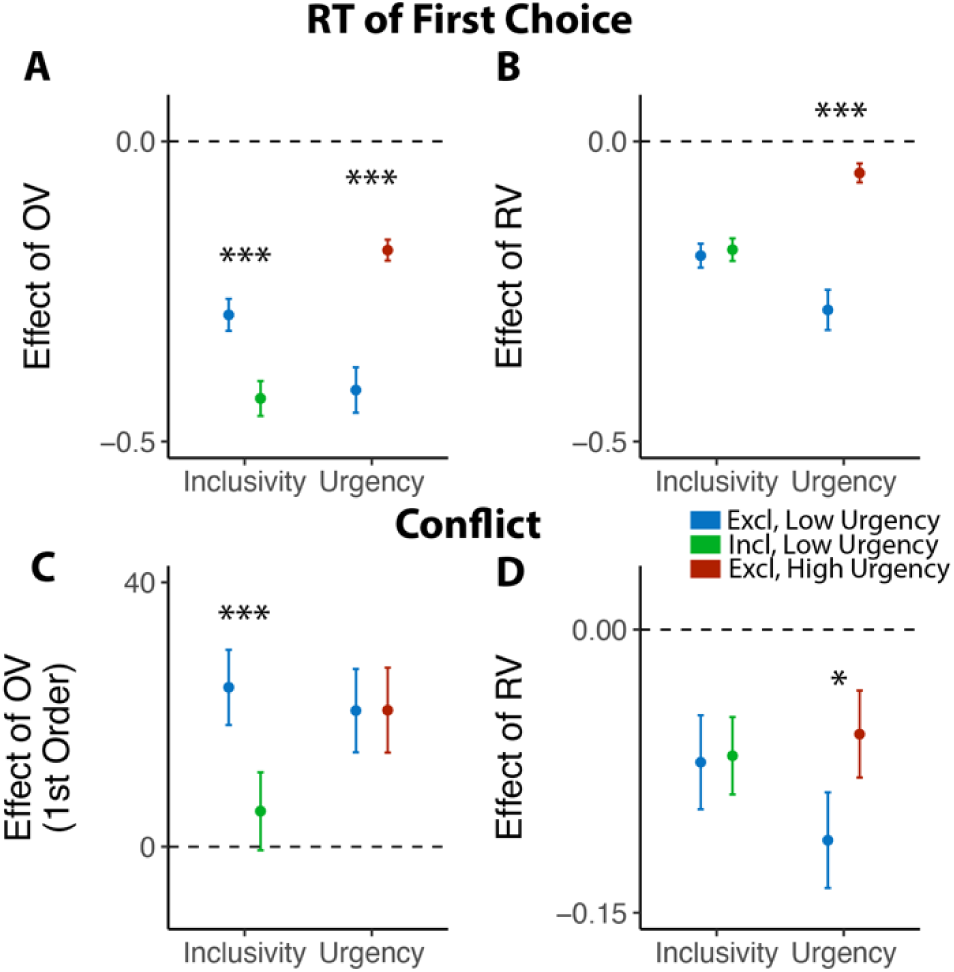
Comparison between the effects of inclusivity (Study 1) and choice urgency (Study 4). In contrast to the effect of choice inclusivity, we found that **(A-B)** choice urgency reduces the effect of overall value and relative value with similar magnitudes; **(C)** choice urgency does not modulate how choice conflict varies with overall value, but **(D)** reduces the negative correlation between relative value and conflict. The error bars indicate standard error. *: p<0.05; ***: p<0.001.

Though inclusivity and urgency can both improve choice behavior, further analyses show that these two methods of choice optimization differ in their ability to improve the subjective experience of choosing. We found that choices with tighter deadlines, despite generating faster choices, did not lead to lower experiences of choice conflict (M_low urgency_=2.56, M_high urgency_=2.58; β *_urgency_*=0.02, 95% CI=[-0.03,0.07], p=0.42). Instead, urgency seems to undercut one of the features of choice sets that typically promotes lower choice conflict: relative value. Across both exclusive and inclusive choices in Study 1 and our low-urgency exclusive choices in Study 4, we found that people experience less conflict the higher the relative value of their choice set (i.e., the easier their choice), consistent with past findings^13,19^. This reduction in choice conflict with relative value was reduced in our high-urgency choices (β*_RV, low urgency_*=-0.11, 95% CI=[-0.16,-0.06], p<0.001; β*_RV, high urgency_*=-0.06, 95% CI=[-0.10,0.01], p=0.016; β*_urgency_*_×*RV*_=0.05, 95% CI = [0.01, 0.09], p=0.021; Figure 7D; see also Figure S10 in Supplementary Materials), while the increase in choice conflict with higher levels of overall value (which was selectively reduced by inclusivity in Study 1) remained unchanged (β*_urgency_*_×*OV_linear*_=-0.38, 95% CI = [-4.66, 3.90], p=0.863; β *_urgency_*_×*OV_quad*_=1.80, 95% CI = [-2.19, 5.79], p=0.377; Figure 7C).

### Choosing how many options to choose

While we have so far focused on comparing inclusive choices to isomorphic choices under an exclusive framing (i.e., by examining only the first choice made in the inclusive condition), these choices afford us a unique opportunity to understand how people evaluate options under conditions where choice is voluntary. In particular, we could examine how people choose how many items to (de-)select, and how this was related to experiences of choice conflict towards the initial set of options. We found that these voluntary choices were heavily determined by the overall value of the choice set, such that participants selected more options (β*_OV_linear_*=49.11, 95% CI=[44.24,53.97], p<0.001; β*_OV_quad_*=6.17, 95% CI=[3.09, 9.26], p<0.001) and removed fewer options (β*_OV_linear_*=-47.59, 95% CI=[-52.31,-42.86], p<0.001; β*_OV_quad_*=10.72, 95% CI=[8.43,13.01], p<0.001) the more valuable those options (Figure 8A). These findings are remarkable given that these choices were entirely optional (i.e., participants could have chosen to move on to the next trial at any point) and none of these products were inherently aversive (i.e., participants could have always chosen all of the options on each trial). These findings are also independent of the influence of relative value, which was negatively correlated with additional option selection (β*_RV_*=-0.06, 95% CI=[-0.08,-0.03], p<0.001) and removal (β*_RV_*=-0.02, 95% CI=[-0.04,-0.00], p=0.014; Figure 8B). These analyses also control for the speed and accuracy of the initial choice one makes from that set. We found that the indifference point of whether each particular item was chosen or removed aligned with the average value of all the items that the individual had assessed - items that exceeded this average were chosen (Studies 1 and 3A) and retained (Studies 2 and 3B), while items that fell below this average were not kept (Figure 8C).

**Figure 8.**
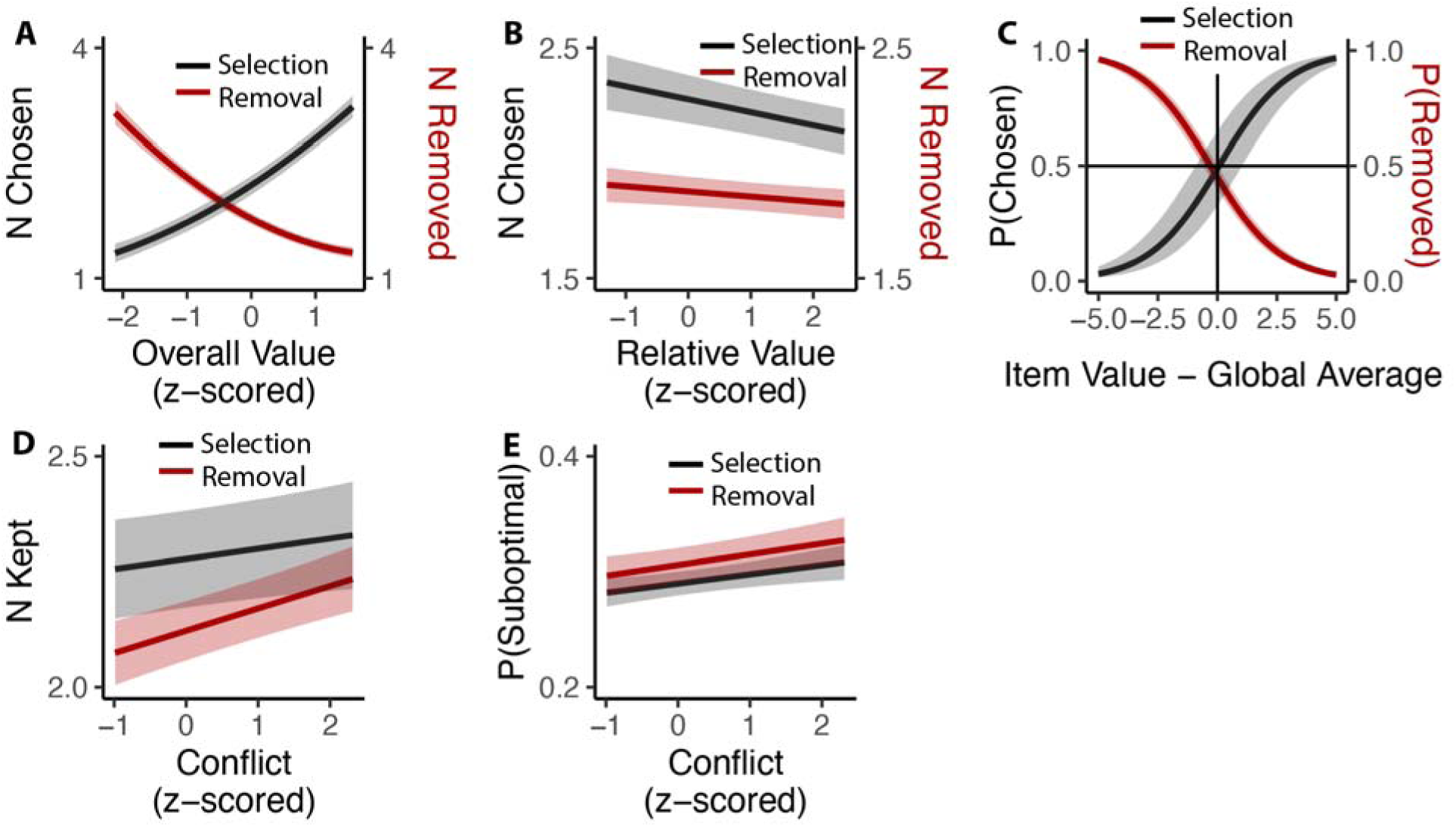
Influences of choice set values on additional inclusive decision-making. **(A)** Influence of overall value on voluntary decision-making. As the overall value increased, participants chose more options in the selection task (Black) and removed fewer options in the removal task (Red). **(B)** Participants made fewer decisions as the relative value increased, regardless of selection or deselection tasks. **(C)** Inclusive choices were guided by item values relative to the global average. Items with value higher than average were more likely to be selected and less likely to be removed. **(D)** Participants kept more options (select more or remove less) when they experienced more conflict. **(E)** This leads to higher likelihood of keeping unfavored options with higher conflict. Shaded areas indicate 95% confidence intervals.

We then examined whether there was a relationship between how conflicted participants reported feeling when faced with the initial set of four options and how many options they ended up choosing on that trial (again, focusing only on inclusive choices). We found that experiencing the initial choice as more conflicting led participants to select more options in Studies 1 and 3A (β*_conflict_*=0.023, 95% CI=[0.001,0.045], p=0.042; Figure 8D) and to remove fewer options (e.g., to keep more options) in Studies 2 and 3B (β*_conflict_*=0.051, 95% CI=[0.033,0.069], p<0.001; Figure 9D). Collapsing across these studies, we saw that higher levels of choice conflict were associated with sequences of decisions that ended with participants keeping a larger number of options (either through acquisition or retention; β*_conflict_*=0.038, 95% C I: [0.024,0.051], p<0.001). To examine whether keeping these additional options on high-conflict trials reflected more or less optimal decision-making, we counted the number of options that were kept on a given trial despite having a value lower than the within-subject mean value of all possible options in the study. Controlling for overall value and relative value, we found that higher levels of experienced choice conflict at the start of a given choice set predicted keeping a higher proportion of these sub-par options (selection: β*_conflict_*=0.007, 95% CI=[0.002, 0.013], p=0.007; removal: β*_conflict_*=0.010, 95% CI=[0.004,0.016], p=0.001; combining selection and removal: β *_conflict_*=0.009, 95% CI=[0.005, 0.013], p<0.001; Figure 8E).

## Discussion

Decision making is at the core of some of the most demanding tasks we face every day, and can create significant bottlenecks to completing those tasks. Humans are vexed by choices large and small because they nearly all produce the same tension: choosing some options means giving up on others. Here, we investigated whether choice behavior and experience were improved by relaxing this tension through greater choice inclusivity. We found that participants were more efficient and less conflicted choosers when making inclusive choices, independent of the choice goal (selection or removal) and in both the presence and absence of time pressure. We showed that the patterns of choice behavior we observed when participants were making inclusive relative to exclusive choices – including a selective enhancement of the influence of overall value on RT without altering the influence of other value estimates on behavior – was uniquely accounted for by a model in which choice inclusivity resulted in a relaxation of mutual inhibition between the competing options. These patterns of behavior and choice conflict were distinct from those resulting from a change in response deadline, suggesting a unique benefit from choice inclusivity compared to urgency-based strategy of choice optimization.

While our studies provide evidence of a task context selectively altering levels of mutual inhibition while holding all the other parameters of decision process constant, this raises the question of whether these alterations are implemented via top-down control or construction of evidence for and against each option. For example, whereas our modeling assumed a form of lateral inhibition between candidate responses, other models have proposed that this inhibition occurs through a feedforward route^41^. This form of feedforward inhibition, whereby positive evidence for one option results in negative evidence for others, could be seen as reflecting the role of opportunity costs (i.e., the value of options foregone)^42^ in the decision process. From this perspective, it is possible to imagine that inclusive choices engender less of a feeling of anticipated loss from not selecting a particular option, because this option will remain available subsequently.

Making choices more inclusive led to an overall reduction in experiences of choice conflict, but this inclusivity benefit varied in important ways with overall value. When participants were selecting which options to obtain (Study 1), inclusivity most benefited the upper arm of the U-shaped curve (high-value choices), presumably because this relieved the tension of not being able to choose more than one of these; conversely, low-value choices engendered a similar level of conflict irrespective of their inclusivity because participants were still constrained by having to choose one of these. Confirming this interpretation, when we instead endowed participants with these options and asked them which ones to remove (Study 2), the interaction between inclusivity and overall value reversed: now, inclusivity selectively benefited choices between low-value options (in which cases participants could opt to remove all of their options) more than choices between high-value options (in which cases participants were now faced with the dilemma of having to drop at least one of these). These findings have important implications for understanding the mechanistic basis for experiences of choice conflict.

Decision-making dysfunctions are common across a wide range of psychiatric disorders^43^, such as generalized anxiety disorder^44^ and obsessive-compulsive disorders^10^. For such individuals, decision-making can be particularly aversive (e.g., anxiety-provoking) and even lead to extreme indecision and choice paralysis, resulting in decisions being prolonged, deferred, or avoided altogether. Our findings point to potential mechanisms contributing to these affective and behavioral sequelae, suggesting that they may stem in part from aberrant levels of competition resulting from excessive levels of mutual inhibition between candidate responses. This in turn suggests directions for follow-up research aimed at better understanding etiology, classification, and treatment for these disorders.

Our findings suggest that real-world choices can be improved by offering inclusivity rather than exclusivity between options. With that said, many real-world choices are by definition exclusive (e.g., requiring payment for each additional option). This places significant limits on the potential for generalizing our findings to applications in the marketplace and elsewhere. Nevertheless, it is interesting to speculate whether similar benefits could accrue in these cases if one considers inclusivity over a longer time horizon (e.g., that they will have the opportunity to purchase other options in the future rather than in the moment). Future research would benefit from examining the limits of inclusivity in its various forms (e.g., convenient returns, tasting menus) on choice, and informing policy accordingly. Future work should also explore the feasibility of inducing inclusivity as an internal mindset rather than external choice conditions. For example, decision-makers can be encouraged to evaluate options in isolation rather than in comparison to one another.

Our computational and empirical findings point to a deeper puzzle: if mutual inhibition is maladaptive for optimizing decisions and experiences thereof, what benefit does it afford? Before examining this further, it is important to note that mutual inhibition’s role in choice does not appear to be universal in the animal world. For instance, starlings make value-based decisions in a manner that resembles a race process (i.e., with limited or no mutual inhibition)^45^, suggesting that mutual inhibition reflects an evolutionary adaptation within the circuits that support decision-making. While identifying this adaptive role is well outside of the scope of what our studies can speak to, we can offer two speculations. First, while mutual inhibition may not be locally adaptive for selecting between responses to value-based decisions as in our experiment, other work has shown that such inhibition may benefit other cognitive processes including monitoring (e.g., detecting levels of conflict to guide control allocation)^46^ and/or separating neuronal representations held in working memory that guide behavior^47,48^, both plausibly processes that are expanded in humans relative to other species. Second, while choice values were not dependent on one another within our choice sets, it is possible that such (inverse) dependencies arise often enough in real-world choice settings that individuals develop priors that approximate mutual inhibition (e.g., assumptions that certain feature values are consistently anti-correlated). This latter possibility can be explored further by examining ecological data and the trajectory of learning through choice environments with varying statistical structure.

In addition to elucidating mechanisms of choice competition in typical one-shot choices, our findings also provide valuable insights into how humans and other animals make decisions in sequential environments, namely under conditions where they can choose when to stop choosing ^49^. We found that these choices were primarily guided by the value of those individual items relative to the global average value of items that person had evaluated, consistent with normative and empirical research on foraging decisions^50^. Interestingly, we found preliminary evidence that the number of options a participant chose was associated with the level of choice conflict they experienced while making the initial choice. In this way, the current work lays the groundwork not only for understanding forms of decision paralysis that occur when having to make a single choice, but also pathological behaviors like over-consumption and hoarding that might occur in contexts where multiple choices are allowed, indeed including the buffet.

## Methods and Materials

### Ethical compliance statement

Across all five studies, participants (N=462) received monetary compensation, and provided informed consent in a manner approved by Brown University’s Institutional Review Board under protocol 1606001529. No statistical methods were used to pre-determine sample sizes but our sample sizes are larger than those reported in previous publications^32^.

### Study 1

#### Participants

17 participants (4 females, 13 males; age = 21±1 ys) participated in Study 1A (in-lab), and 74 (35 females, 39 males; age = 36±10 ys) participants were recruited for Study 1B, an online replication study on Prolific. Participants were excluded from our analysis based on the following criteria: (1) to ensure that participants’ product ratings prior to the choice task cover the full range of the liking scale, we excluded participants whose standard deviation of their product ratings was too low (*SD_value_*<1) or too high (*SD_value_*>5); (2) to ensure compliance with the task instructions, we calculated participants’ choice consistency within the easy trials (defined as trials with relative value greater than the within-participant median), and excluded participants whose mean accuracy in easy trials was less than 25%; (3) we also excluded participants with too low variance in their conflict ratings (*SD_conflict_*<0.5). This resulted in a sample of 65 participants for Study 1B (30 females, 35 males; age = 37±11 ys) and no exclusions for Study 1A. The qualitative patterns reported in this paper hold when we include all 74 Study 1B participants.

#### Procedure

Our experiment consisted of three phases (Figure 1). In Phase 1, participants viewed a series of products (in-person: 359, online: 200) and were instructed to rate how much they would like to have each one, by clicking on an analog liking scale from 0 (not at all) to 10 (a lot). In Phase 2, participants made choices (in-person: 160, online: 120) among sets of four products. For each set, we sample from a uniform distribution of overall value ([0,10]), and then sample from the distribution of possible best value given the sampled overall value ([OV, min(10,4*OV)]). Then we can calculate the mean of the remaining options, and then generate the second-best option from all possible alternatives. We repeat this until all options are generated for that trial. This process is performed separately for inclusive and exclusive conditions, with distributions of overall value and relative value matched between these two conditions. In addition, we followed two constraints: 1) for each product, it will be displayed at most 3 times; 2) for each product, it will not be displayed for two consecutive trials; and 3) across all trials, there will not be two sets of alternatives with the same products. On *exclusive* choice trials, participants were allowed to choose one product from the choice set. Once they clicked on this product, a box appeared around it and they proceeded to the next trial. On *inclusive* choice trials, participants were able to continue selecting as many options as they preferred after they chose the first one. The two choice conditions were intermixed, occurred with equal likelihood, and were cued by the color of the fixation cross in the middle of the screen (blue for exclusive choices and green for inclusive choices). In both conditions, participants were given up to 9s to complete each trial and, importantly, were instructed to always start by selecting their favorite option out of the set. In Phase 3, participants viewed each choice set again and rated the level of conflict they felt when facing each set on a 5-point scale.

#### Analysis of choice behavior

For the choice phase, we used linear mixed effect regressions (R package lme4) ^51^ to analyze reaction time (RT) and generalized linear mixed effect regression with logistic transformation for choice accuracy (whether the highest-rated option was selected) for the first choice in each condition. All regressions include choice inclusivity (coded with successive differences contrast so that intercept is the average across two conditions and the contrast is the difference between two conditions)^52^, the overall (mean) value of the choice set, the relative value (quantified as the difference between the value of the highest-rated product and the mean value of the remaining products), the interactions between choice inclusivity and overall/relative value, and trial order, with random (subject-specific) intercept and slopes for each variable ^53^.

#### LCA simulation

We modified the Leaky Competing Accumulator model (LCA; Figure 3A)^34,35^ to simulate choice behavior. In this LCA model, option-specific leaky accumulators accumulate evidence until one of the accumulators reaches a decision boundary (starting at *a* and collapsing at the rate of *ϑ*) and induces a response. The first boundary-crossing time and the corresponding option are recorded as the response time and the choice. At each time step, the accumulation process advances as

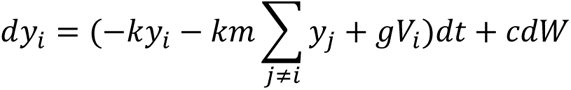

where *V_i_* is the input from option *i* in the choice set, *g* is the gain of input, *k* denotes the decay of the leaky accumulator, *m* represents the ratio between mutual inhibition from other accumulators and decay, and *cdW* is the Gaussian random noise with mean 0 and variance *c*.

We first fixed *g*, *c* and manipulated parameters *k*, *m*, *a*, *ϑ*. We simulated the choice behavior (reaction time and accuracy of the first choice; 100 iterations per combination of parameters) for different combinations of option values across a range of these four parameters. We then performed the same linear and generalized linear regressions on these simulated data as for the empirical data (e.g., regressing simulated RT and accuracy on overall value and relative value) to compare those findings **qualitatively** with those observed across our experimental conditions. We then performed the same process with varying *g* to confirm that the observed qualitative pattern is consistent across different levels of input gain.

To confirm that manipulation of *m* can generate observed behavioral patterns in the empirical data, we performed a grid search across different combinations of *k*, *g*, *a*, *ϑ* with high and low levels of *m* (representing exclusive and inclusive conditions), and identified the best parameter set that maximizes the similarity between simulated and empirical regression estimates. We then compare the simulated regression estimates with empirical ones to confirm that manipulating *m* can generate the observed pattern (Figure 3D). To compare the manipulation of *m* with the tuning of decision boundary parameters (*a* and *ϑ*), we performed additional grid search across different combinations of *k*, *g*, *m* with varying *a* and *ϑ* (for exclusive and inclusive conditions), and compared the best predictions with the outcome from manipulations of *m* (Figure S3 in Supplementary Materiels).

#### Analysis of choice conflict

For the conflict rating phase, we used linear mixed-effect regressions to analyze the rating of choice conflict. All regressions include choice inclusivity, the linear and quadratic terms (using orthogonal polynomials) of the overall (mean) value of the choice set, the relative value (quantified as the difference between the value of the highest-rated product and the mean value of the remaining products), the interactions between choice inclusivity and overall/relative values, and trial order, with random (subject-specific) intercept and slopes for each variable. We also tested additional models with control of reaction time and choice accuracy (see Supplementary Materials).

### Study 2

#### Participants

118 participants (59 females, 59 males; age = 36±11 ys) participated were recruited for this study on Prolific. Participants were excluded from our analysis based on the same criteria for Study 1. This resulted in a sample of 98 participants for Study 2 (50 females, 48 males; age = 35±11 ys). The qualitative patterns reported in this paper hold when we include all 118 participants.

#### Procedure

The procedure of the experiment is the same with Study 1, except that participants were instructed to make deselections among sets of four pre-selected products framed with boxes. On *exclusive* choice trials, participants were allowed to deselect one product from the choice set. Once they clicked on this product, the box around it disappeared and they proceeded to the next trial. On *inclusive* choice trials, participants were able to continue deselecting as many options as they preferred after they deselected the first one. The two choice conditions were intermixed, occurred with equal likelihood, and were cued by the color of the fixation cross in the middle of the screen (blue for exclusive deselections and green for inclusive deselections). In both conditions, participants were given up to 9s to complete each trial and, importantly, were instructed to always start by deselecting their least favorite option out of the set. In Phase 3, participants viewed each choice set again and rated the amount of conflict they felt when facing each set on a 5-point scale.

#### Analysis

For the choice phase, we analyzed reaction time (RT) and choice accuracy (whether the lowest-rated option was deselected) for the first choice in each condition. The setup of predictors is the same with Study 1, except for the relative value (quantified as the absolute difference between the value of the lowest-rated product and the mean value of the remaining products).

### Study 3

#### Participants

68 participants (21 females, 47 males; age = 33±8 ys) participated in the selection task (Study 3A), and 78 (40 females, 38 males; age = 34±10 ys) participants were recruited for the deselection task (Study 3B). Participants were excluded from our analysis based on the same criteria for Study 1 and 2. This resulted in a sample of 59 participants for selection task (19 females, 40 males; age = 33±8 ys) and 61 (32 females; 29 males; age = 34±9 ys) for the deselection task. The qualitative patterns reported in this paper hold when we include all participants.

#### Procedure

The replication studies follow the same procedures of selection (Study 1) and deselection study (Study 2) with the variance that 1) we removed the choice deadline and 2) we selected a new set of products (N=210).

#### Analysis

We followed the same analysis settings in Study 1-2. The only difference is that we log-transformed the reaction time to account for the long-tail distribution.

### Study 4

#### Participants

108 participants (50 females, 58 males; age = 36±9 ys) participated were recruited for this study on Prolific. In addition to the exclusion criteria for Study 1, participants with high omission rate in high urgency choice trials (>=40%) were also excluded from our analysis. This resulted in a sample of 85 participants for Study 4 (39 females, 46 males; age = 36±9 ys). The qualitative patterns reported in this paper hold when we include all 108 participants.

#### Procedure

The procedure of the experiment is the same with Study 1, except that participants were instructed to make *exclusive* selections with low or high urgency. On *low urgency* choice trials, participants have unlimited time to make their choice. On *high urgency* choice trials, participants have only **3 seconds** to make their choice. The two choice conditions were intermixed, occurred with equal likelihood, and were cued by the color of the fixation cross in the middle of the screen (blue for low urgency choice trials and green for high urgency choice trials). In Phase 3, participants viewed each choice set again and rated the amount of conflict they felt when facing each set on a 5-point scale.

#### Analysis

The setup of analysis is the same with Study 1, except now low and high urgency are coded as successive difference contrast in the model.

### Analysis of inclusive choices

We only included inclusive choice in this analysis and combined studies based on type of choice (selection: Study 1 and Study 3A; removal: Study 2 and Study 3B). We analyzed the number of inclusive choices in each trial by fitting linear mixed models to **(a)** number of selections/removals and **(b)** number of products kept for each trial (for removal study, it refers the size of choice set minus number of choices). We then examined the quality of subsequent choices. We analyzed **(c)** probability of selection/removal per product and **(d)** likelihood of keeping suboptimal products (with value lower than global average) per trial. For (a), (b) and (d), the model included the linear and quadratic terms (using orthogonal polynomials) of the overall (mean) value of the choice set, the relative value (quantified as the difference between the value of the highest-rated product and the mean value of the remaining products), the reaction time and accuracy of initial choices, and trial order. For (c), the model included overall value and the difference between product value and subject-specific global average. For (b) and (d), we also performed the same analysis after combining selection and removal studies. All of these models included random (subject-specific) intercept and slopes for each variable.

### Data availability

All experiment de-identified data is publicly available at https://github.com/Jasonleng/choiceinclusivity.git.

This will be shared upon publication.

### Code availability

Data analysis script notebooks and simulation code are publicly available at https://github.com/Jasonleng/choiceinclusivity.git.

This will be shared upon publication.

## Acknowledgments

This work was funded by a Center of Biomedical Research Excellence grant P20GM103645 from the National Institute of General Medical Sciences (A.S.), grant R01MH124849 from the National Institute of Mental Health (A.S.). The authors are grateful to Gloria Feng for assistance in data collection, Harrison Ritz for generously sharing code and advice, and to Matthew R. Nassar, Michael J. Frank, Frederick Callaway, Alexander Fengler, Yi-Hsin Su and Joonhwa Kim for helpful discussion.

## Author Contributions Statement

X. L., R. F., and A. S. developed the idea and planned the original study. R. F. and A. S. supervised the project. X. L., R. F., and T. S. implemented the experiment. X. L. and R. F. performed the data analysis and computational modeling. X. L. wrote the initial draft. X. L., R. F., and A. S. revised the manuscript.

## Competing Interests Statement

The authors have no competing interests.

## Notes

### Competing Interest Statement

The authors have declared no competing interest.

### Summary of Updates

Updated figure and included additional data.

