## Supplementary Materials for "Mutual inclusivity improves decision-making by smoothing out choice’s competitive edge"

### 1. Study S1 - Initial and subsequent choices temporally separated for inclusive choices

Our experiments allow people to make all the choices together for each trial in the inclusive condition. It could be that the observed influence of choice inclusivity benefited from the fact that people can make their subsequent choice immediately after their first choice so that it is natural for them to accumulate evidence for each option independently. To confirm that our findings do not vary with the temporal distance between initial and subsequent choices, we performed an additional Study (N = 77) in which inclusive choices are separated into two phases. Choice trials are divided into mini-blocks in which people will only perform one type of choice (inclusive or exclusive). In the inclusive blocks, people will experience all trials and make their first choices. After that, they can revisit these trials and choose from the remaining ones. We followed the same analysis in Study 1 and 3A. Our findings in this supplementary study are consistent with the patterns in Study 1 and Study 3A, suggesting that the influence of inclusivity is not a side-product of being able to complete choices together.

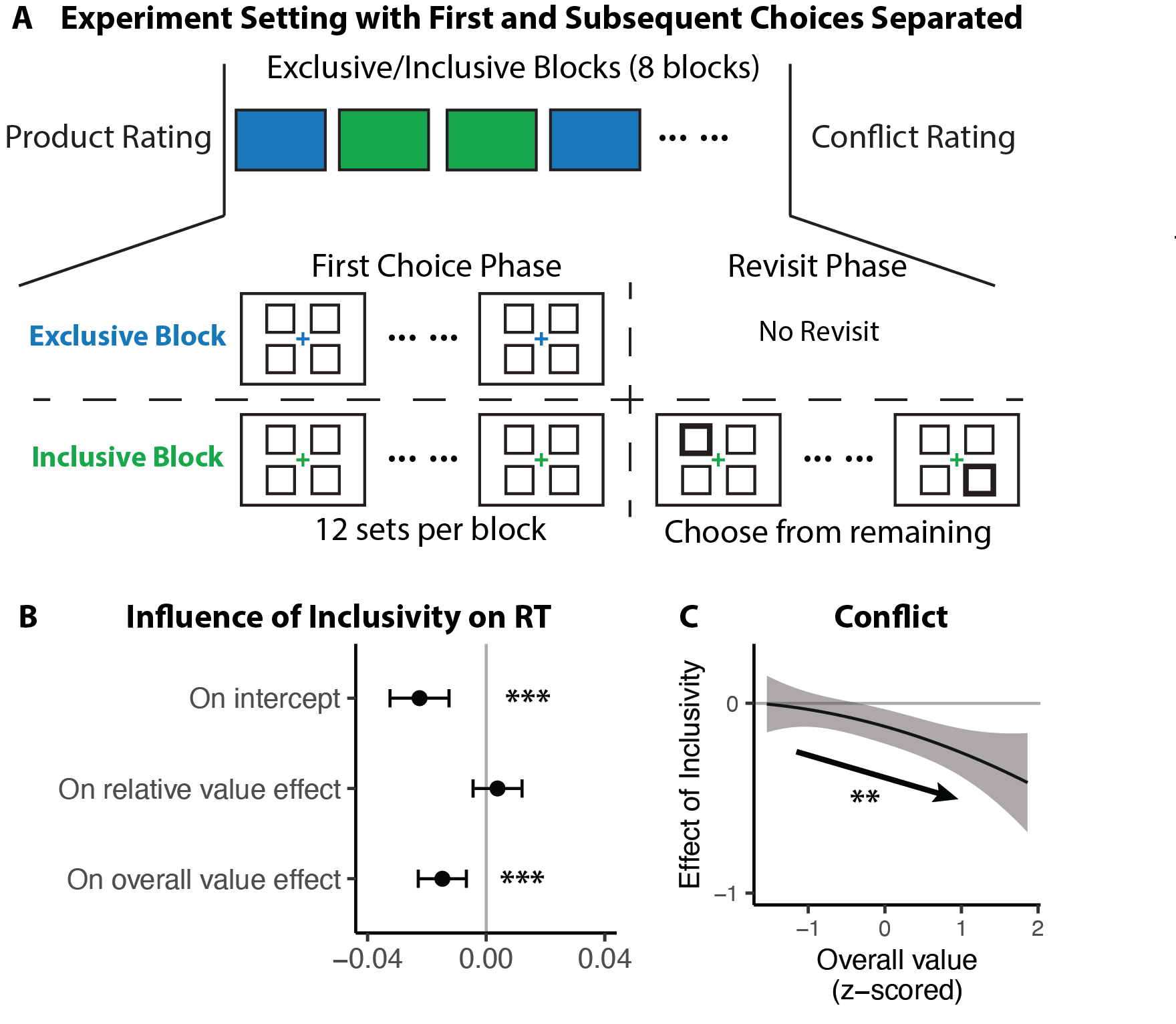

**Figure S1. Complementary study with initial and subsequent choices separated for inclusive choices. A)** Experiment setting. The choice part is separated into mini-blocks, either inclusive or exclusive. In inclusive blocks, participants first chose one item from each trial and revisited these trials at the end of the block to make subsequent choices if they wanted. **B)** The influence of inclusivity on the RT of first choices replicates the findings in Study 1. **C)** Consistent with Study 1 and Study 3A, participants experience less conflict when they can revisit and choose later, especially when the overall value of the choice set is high. Error bars and shades represent 95% confidence intervals. **: p<0.01; ***: p<0.001.

### 2. Subset of Study S1 with incentive-compatible setting

It is possible that these findings rely on the hypothetical choice setting in the study. To confirm that our findings persist in an incentive-compatible setting, we introduced incentives to a subset of participants in Study S1 (N=38). We framed the experiment as a gift raffle game in which participants would receive a real gift from the pool of products. Before the start of the experiment, participants were told that they could receive a gift and they were instructed to provide information about their preferred methods of gift delivery. Before making choices, the probability of getting each item is the same and low. We informed participants that their choices would increase their chances of winning the chosen item. After the experiment, participants will have a chance to receive a product randomly drawn based on the probability distribution modified by participants’ choices.

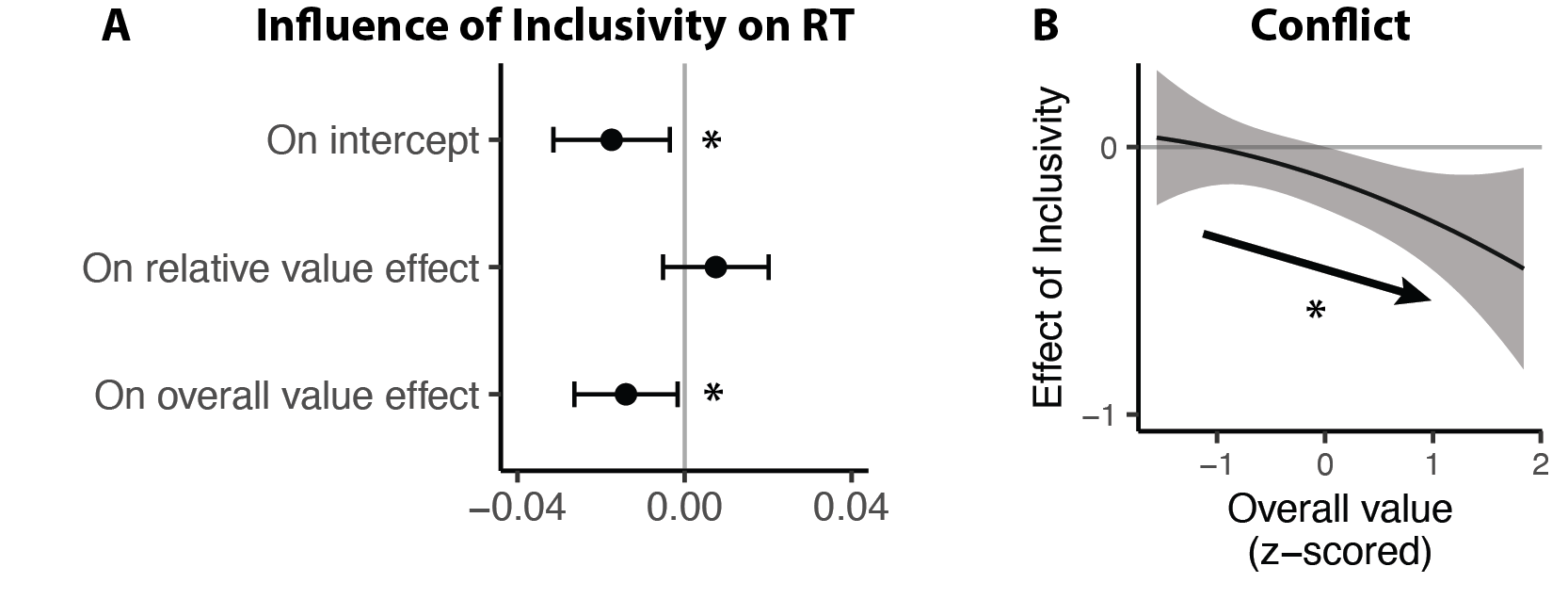

**Figure S2. A subset of Study S1 with compatible incentive.** In a subset of Study S1, participants will have a chance to receive a gift with probability aligned with their choices. A) The influence of inclusivity on the RT of first choices replicates the findings in Study 1 and Study 3A. B) Consistent with Study 1 and Study 3A, participants experience less conflict when they can revisit and choose later, especially when the overall value of the choice set is high. Error bars and shades represent 95% confidence intervals. *: p<0.05.

### 3. Comparison between competition-varying and boundary-varying LCA model simulation

We further confirm that varying competition but not boundary parameters in the LCA model is essential for capturing the observed behavior pattern. We first run three additional simulations in which competition is the same across conditions while the initial threshold and/or boundary collapse rate varies between conditions. Consistent with our predictions, we found that these simulations cannot capture the observed pattern, especially the selective influence of inclusivity on the speeding effect of OV. To validate that competition is essential even if the model enables free adjustment of all three parameters (competition, initial threshold, boundary collapse rate), we ran another simulation with all three parameters freely varying between the two conditions. We then inspected the change of these parameters in the best simulation settings. We found that competition still varies strongly between these two conditions. Importantly, we did not see a large adjustment of the initial threshold.

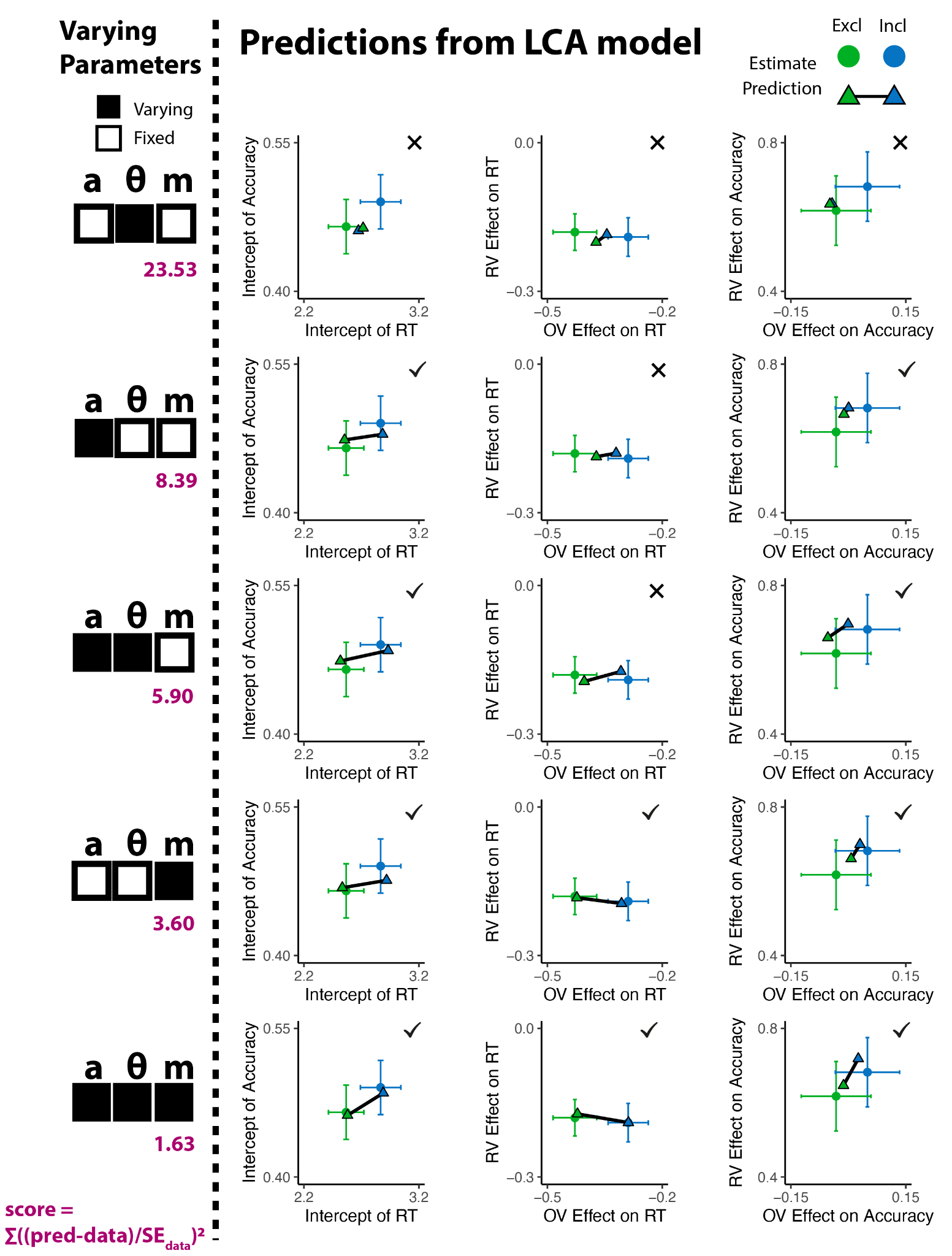

**Figure S3. Predictions from the LCA model with varying boundary parameters.** Models without varying mutual inhibition m (top three rows) cannot predict the observed relationship between inclusivity and the speeding effects of OV/RV. Models with varying mutual inhibition m (bottom two rows) can generate predictions matched with empirical findings. Error bars indicate 95% confidence intervals for estimated parameters.

**Table S1 Parameter settings for LCA model with varying theta, a, and m**

|  | Inclusive | Exclusive |
| --- | --- | --- |
| **competition m** | **0.2** | **1.6** |
| Threshold a | 0.9 | 0.9 |
| theta/rad | 0.18 | 0.23 |

### 4.  Mixed-effect model results for Study 1

**Table S2. Mixed-effect models of first choice reaction time and accuracy in Study 1**

|  | **Reaction Time** | | | **Accuracy** | | |
| --- | --- | --- | --- | --- | --- | --- |
| ***Predictors*** | ***Estimates*** | ***CI*** | ***p*** | ***Log-Odds*** | ***CI*** | ***p*** |
| Intercept | 2.72 | 2.56 – 2.88 | **<0.001** | -0.09 | -0.19 – 0.01 | *0.083* |
| Inclusivity | -0.30 | -0.38 – -0.22 | **<0.001** | -0.10 | -0.19 – -0.01 | **0.029** |
| Overall Value | -0.36 | -0.41 – -0.31 | **<0.001** | 0.01 | -0.06 – 0.08 | 0.791 |
| Relative Value | -0.19 | -0.22 – -0.16 | **<0.001** | 0.65 | 0.57 – 0.73 | **<0.001** |
| Trial Order | -0.12 | -0.16 – -0.08 | **<0.001** | -0.03 | -0.08 – 0.01 | 0.164 |
| Incl by OV | -0.14 | -0.20 – -0.08 | **<0.001** | -0.08 | -0.19 – 0.03 | 0.148 |
| Incl by RV | 0.01 | -0.04 – 0.06 | 0.666 | -0.06 | -0.17 – 0.04 | 0.246 |

**Table S3. Mixed-effect models of conflict rating in Study 1**

|  | **Conflict** | | |
| --- | --- | --- | --- |
| ***Predictors*** | ***Estimates*** | ***CI*** | ***p*** |
| Intercept | 2.38 | 2.25 – 2.51 | **<0.001** |
| Inclusivity | -0.46 | -0.62 – -0.29 | **<0.001** |
| Overall Value | 14.71 | 4.10 – 25.32 | **0.007** |
| Overall Value^2 | 12.66 | 7.62 – 17.70 | **<0.001** |
| Relative Value | -0.07 | -0.11 – -0.03 | **<0.001** |
| Trial Order | 0.01 | -0.03 – 0.05 | 0.716 |
| Incl by OV | -18.76 | -27.07 – -10.45 | **<0.001** |
| Incl by OV^2 | -4.87 | -9.97 – 0.23 | 0.061 |
| Incl by RV | 0.00 | -0.05 – 0.05 | 0.894 |

**Table S4. Mixed-effect model of conflict rating in Study 1 controlling for RT and accuracy**

|  | **Conflict** | | |
| --- | --- | --- | --- |
| ***Predictors*** | ***Estimates*** | ***CI*** | ***p*** |
| Intercept | 2.36 | 2.23 – 2.50 | <0.001 |
| RT | 0.09 | 0.05 – 0.13 | <0.001 |
| Accuracy | -0.09 | -0.13 – -0.05 | <0.001 |
| Inclusivity | -0.44 | -0.59 – -0.28 | <0.001 |
| OV | 19.55 | 10.17 – 28.93 | <0.001 |
| OV^2 | 11.11 | 6.56 – 15.66 | <0.001 |
| RV | -0.03 | -0.06 – 0.00 | 0.075 |
| Trial Num | 0.01 | -0.03 – 0.05 | 0.696 |
| Incl by OV | -17.43 | -25.30 – -9.56 | <0.001 |
| Incl by OV^2 | -5.03 | -10.05 – -0.02 | 0.049 |
| Excl by RV | -0.01 | -0.06 – 0.05 | 0.836 |

### 5. Mixed-effect model results for Study 2

**Table S5. Mixed-effect models of first choice reaction time and accuracy in Study 2**

|  | **Reaction Time** | | | **Accuracy** | | |
| --- | --- | --- | --- | --- | --- | --- |
| ***Predictors*** | ***Estimates*** | ***CI*** | ***p*** | ***Log-Odds*** | ***CI*** | ***p*** |
| Intercept | 3.04 | 2.90 – 3.18 | **<0.001** | -0.25 | -0.34 – -0.16 | **<0.001** |
| Inclusivity | -0.13 | -0.19 – -0.07 | **<0.001** | -0.14 | -0.23 – -0.06 | **0.001** |
| Overall Value | 0.43 | 0.38 – 0.47 | **<0.001** | 0.09 | 0.04 – 0.14 | **0.001** |
| Relative Value | -0.16 | -0.19 – -0.13 | **<0.001** | 0.48 | 0.41 – 0.55 | **<0.001** |
| Trial Order | -0.16 | -0.20 – -0.12 | **<0.001** | -0.05 | -0.09 – -0.01 | **0.023** |
| Incl by OV | 0.07 | 0.02 – 0.12 | **0.003** | 0.06 | -0.03 – 0.15 | 0.165 |
| Incl by RV | -0.01 | -0.05 – 0.04 | 0.772 | -0.08 | -0.17 – 0.01 | 0.070 |

**Table S6. Mixed-effect model of conflict rating in Study 2**

|  | **Conflict** | | |
| --- | --- | --- | --- |
| ***Predictors*** | ***Estimates*** | ***CI*** | ***p*** |
| Intercept | 2.21 | 2.12 – 2.30 | **<0.001** |
| Inclusivity | -0.08 | -0.15 – -0.01 | **0.030** |
| Overall Value | 46.04 | 37.11 – 54.97 | **<0.001** |
| Overall Value^2 | 20.84 | 16.61 – 25.06 | **<0.001** |
| Relative Value | -0.07 | -0.10 – -0.03 | **<0.001** |
| Trial Order | -0.00 | -0.03 – 0.03 | 0.951 |
| Incl by OV | 7.23 | 1.23 – 13.24 | **0.018** |
| Incl by OV^2 | -2.45 | -7.29 – 2.39 | 0.321 |
| Excl by RV | 0.03 | -0.01 – 0.07 | 0.196 |

**Table S7. Mixed-effect model of conflict rating in Study 2 controlling for RT and accuracy**

|  | **Conflict** | | |
| --- | --- | --- | --- |
| ***Predictors*** | ***Estimates*** | ***CI*** | ***p*** |
| Intercept | 2.17 | 2.08 – 2.27 | **<0.001** |
| RT | 0.13 | 0.09 – 0.16 | **<0.001** |
| Accuracy | -0.00 | -0.04 – 0.03 | 0.837 |
| Inclusivity | -0.08 | -0.15 – -0.01 | **0.033** |
| OV | 39.80 | 31.54 – 48.07 | **<0.001** |
| OV^2 | 18.96 | 14.98 – 22.93 | **<0.001** |
| RV | -0.05 | -0.07 – -0.02 | **0.001** |
| Trial Num | 0.01 | -0.03 – 0.04 | 0.695 |
| Incl by OV | 4.40 | -1.54 – 10.34 | 0.147 |
| Incl by OV^2 | -3.34 | -7.97 – 1.29 | 0.158 |
| Excl by RV | 0.03 | -0.01 – 0.07 | 0.170 |

### 6. Distribution of overall and relative values in Study 1 and influence of inclusivity on conflict for low-RV trials.

To confirm that the observed U-shape relationship between overall value and conflict does not evolve from the fact that high- and low-OV trials are also low RV trials, we ran additional analysis which only includes trials with low RV (RV<1). The findings from this analysis are consistent with the mixed-effect model including all trials.

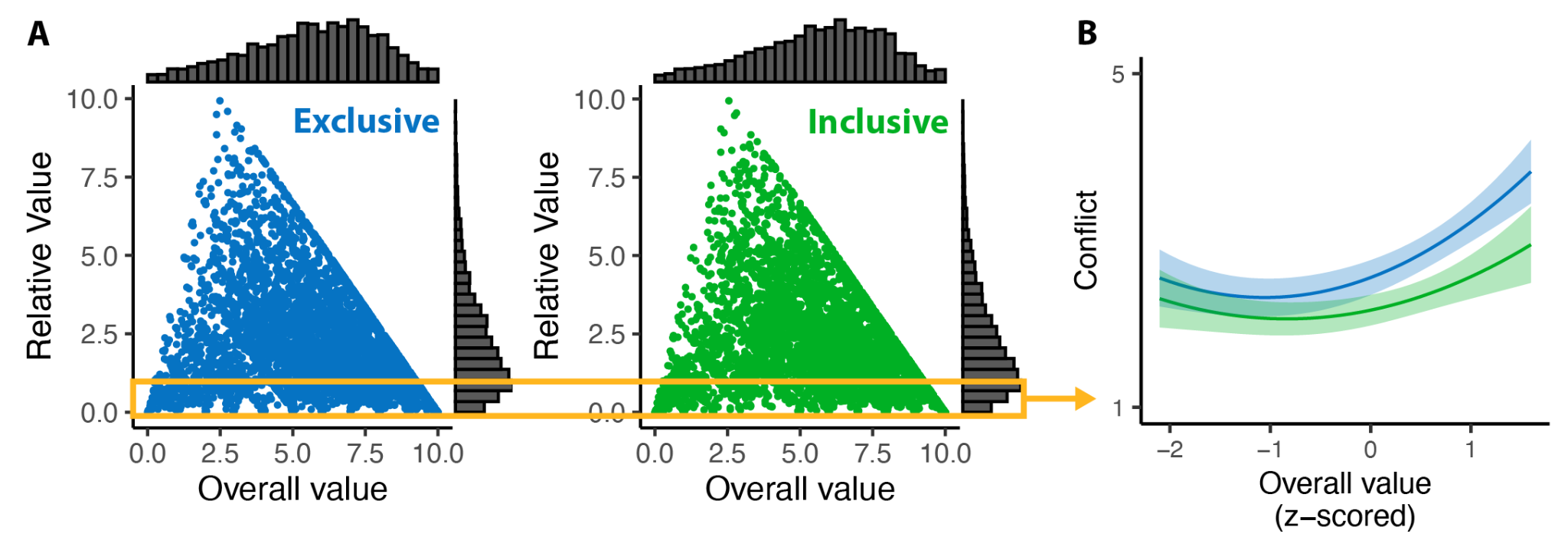

**Figure S4.** A) Distribution of overall value and relative value separated by conditions in Study 1 and B) influence of inclusivity on conflict with varying overall value for trials with low relative value. The orange box in (A) indicates trials included in (B).

### 7. Mixed-effect model results for conflict in Study 1 and Study 3A including only trials where people only chose one.

**Table S8. Mixed-effect model of conflict in Study 1 and Study 3A with only one-choice trials**

|  | **Study 1** | | | **Study 3A** | | |
| --- | --- | --- | --- | --- | --- | --- |
| *Predictors* | *Estimates* | *CI* | *p* | *Estimates* | *CI* | *p* |
| Intercept | 2.35 | 2.20 – 2.50 | **<0.001** | 2.48 | 2.32 – 2.65 | **<0.001** |
| Inclusivity | -0.46 | -0.61 – -0.30 | **<0.001** | -0.29 | -0.45 – -0.13 | **<0.001** |
| Overall Value | 10.38 | 1.88 – 18.87 | **0.017** | -5.36 | -16.17 – 5.46 | 0.332 |
| Overall Value^2 | 8.08 | 3.50 – 12.65 | **0.001** | 7.17 | 2.09 – 12.24 | **0.006** |
| Relative Value | -0.09 | -0.13 – -0.05 | **<0.001** | -0.13 | -0.18 – -0.08 | **<0.001** |
| Trial Order | 0.03 | -0.02 – 0.07 | 0.297 | 0.02 | -0.03 – 0.06 | 0.474 |
| Incl by OV | -19.26 | -28.63 – -9.89 | **<0.001** | -14.03 | -22.35 – -5.71 | **0.001** |
| Incl by OV^2 | -7.24 | -13.88 – -0.60 | **0.033** | -9.68 | -17.28 – -2.08 | **0.013** |
| Incl by RV | -0.04 | -0.10 – 0.03 | 0.242 | -0.06 | -0.15 – 0.02 | 0.139 |

### 8. Mixed-effect model results for Study 3

**Table S9. Mixed-effect model of log-transformed RT and accuracy in Study 3A**

|  | **Replication of Selection Study log-transformed RT** | | | **Replication of Selection Study**  **Accuracy** | | |
| --- | --- | --- | --- | --- | --- | --- |
| *Predictors* | *Estimates* | *CI* | *p* | *Odds Ratios* | *CI* | *p* |
| Intercept | 0.46 | 0.42 – 0.50 | **<0.001** | 0.94 | 0.84 – 1.04 | 0.229 |
| Inclusivity | -0.02 | -0.03 – -0.00 | **0.019** | 1.00 | 0.90 – 1.11 | 0.972 |
| Overall Value | -0.06 | -0.07 – -0.05 | **<0.001** | 1.08 | 1.01 – 1.15 | **0.016** |
| Relative Value | -0.03 | -0.04 – -0.03 | **<0.001** | 1.83 | 1.68 – 1.98 | **<0.001** |
| Trial Order | -0.03 | -0.04 – -0.02 | **<0.001** | 0.96 | 0.91 – 1.02 | 0.182 |
| Incl by OV | -0.02 | -0.03 – -0.01 | **0.001** | 1.03 | 0.92 – 1.15 | 0.587 |
| Incl by RV | -0.00 | -0.01 – 0.00 | 0.389 | 0.89 | 0.79 – 1.01 | 0.067 |

**Table S10. Mixed-effect model of log-transformed RT and accuracy in Study 3B**

|  | **Replication of Removal Study log-transformed RT** | | | **Replication of Removal Study Accuracy** | | |
| --- | --- | --- | --- | --- | --- | --- |
| *Predictors* | *Estimates* | *CI* | *p* | *Odds Ratios* | *CI* | *p* |
| Intercept | 0.45 | 0.41 – 0.49 | **<0.001** | 0.94 | 0.85 – 1.05 | 0.268 |
| Inclusivity | -0.02 | -0.03 – -0.00 | **0.008** | 0.95 | 0.86 – 1.06 | 0.339 |
| Overall Value | 0.07 | 0.06 – 0.08 | **<0.001** | 1.09 | 1.01 – 1.18 | **0.033** |
| Relative Value | -0.03 | -0.03 – -0.02 | **<0.001** | 1.86 | 1.72 – 2.02 | **<0.001** |
| Trial Order | -0.03 | -0.04 – -0.02 | **<0.001** | 0.98 | 0.93 – 1.04 | 0.523 |
| Incl by OV | 0.02 | 0.01 – 0.03 | **<0.001** | 1.11 | 0.99 – 1.24 | 0.075 |
| Incl by RV | -0.00 | -0.01 – 0.01 | 0.801 | 1.04 | 0.91 – 1.18 | 0.588 |

**Table S11. Mixed-effect model of un-transformed RT in Study 3**

|  | **Replication Study-Selection RT** | | | **RT Replication Study-Removal RT** | | |
| --- | --- | --- | --- | --- | --- | --- |
| *Predictors* | *Estimates* | *CI* | *p* | *Estimates* | *CI* | *p* |
| Intercept | 3.38 | 3.09 – 3.68 | **<0.001** | 3.35 | 3.08 – 3.61 | **<0.001** |
| Inclusivity | -0.14 | -0.25 – -0.03 | **0.010** | -0.10 | -0.21 – -0.00 | **0.044** |
| Overall Value | -0.49 | -0.59 – -0.39 | **<0.001** | 0.51 | 0.44 – 0.58 | **<0.001** |
| Relative Value | -0.28 | -0.35 – -0.21 | **<0.001** | -0.26 | -0.32 – -0.20 | **<0.001** |
| Trial Order | -0.25 | -0.32 – -0.19 | **<0.001** | -0.24 | -0.31 – -0.18 | **<0.001** |
| Incl by OV | -0.15 | -0.25 – -0.05 | **0.004** | 0.13 | 0.04 – 0.23 | **0.006** |
| Incl by RV | -0.02 | -0.10 – 0.07 | 0.731 | -0.02 | -0.10 – 0.07 | 0.712 |

**Table S12. Mixed-effect model of conflict in Study 3**

|  | **Replication of Selection Study Conflict** | | | **Replication of Removal Study Conflict** | | |
| --- | --- | --- | --- | --- | --- | --- |
| *Predictors* | *Estimates* | *CI* | *p* | *Estimates* | *CI* | *p* |
| Intercept | 2.47 | 2.32 – 2.62 | **<0.001** | 2.34 | 2.21 – 2.46 | **<0.001** |
| Inclusivity | -0.33 | -0.49 – -0.18 | **<0.001** | -0.08 | -0.18 – 0.03 | 0.176 |
| Overall Value | -2.85 | -16.16 – 10.47 | 0.675 | 50.32 | 42.26 – 58.37 | **<0.001** |
| Overall Value^2 | 13.39 | 8.12 – 18.67 | **<0.001** | 16.22 | 12.25 – 20.20 | **<0.001** |
| Relative Value | -0.09 | -0.13 – -0.05 | **<0.001** | -0.08 | -0.11 – -0.04 | **<0.001** |
| Trial Order | 0.00 | -0.04 – 0.04 | 0.966 | 0.00 | -0.03 – 0.03 | 0.955 |
| Incl by OV | -13.94 | -22.28 – -5.60 | **0.001** | 6.03 | 1.30 – 10.75 | **0.013** |
| Incl by OV^2 | -3.95 | -9.98 – 2.09 | 0.200 | 0.07 | -4.97 – 5.12 | 0.977 |
| Incl by RV | -0.00 | -0.07 – 0.06 | 0.930 | -0.02 | -0.07 – 0.03 | 0.404 |

### 9. Summary of RT by the order of choices in inclusive condition (Study 1)

**Table S13. Mean of RTs by the order of choices separated by number of chosen options**

| **# of Chosen** | **1st RT** | **2nd RT** | **3rd RT** | **4th RT** | **Total Time (s)** | **# of Trials** |
| --- | --- | --- | --- | --- | --- | --- |
| **1** | 2.82 |  |  |  | 2.82 | 1655 |
| **2** | 2.60 | 1.26 |  |  | 3.86 | 1395 |
| **3** | 2.37 | 1.08 | 1.01 |  | 4.47 | 1040 |
| **4** | 2.18 | 0.99 | 0.84 | 0.70 | 4.71 | 980 |

### 10. Supplementary figures for Study 3

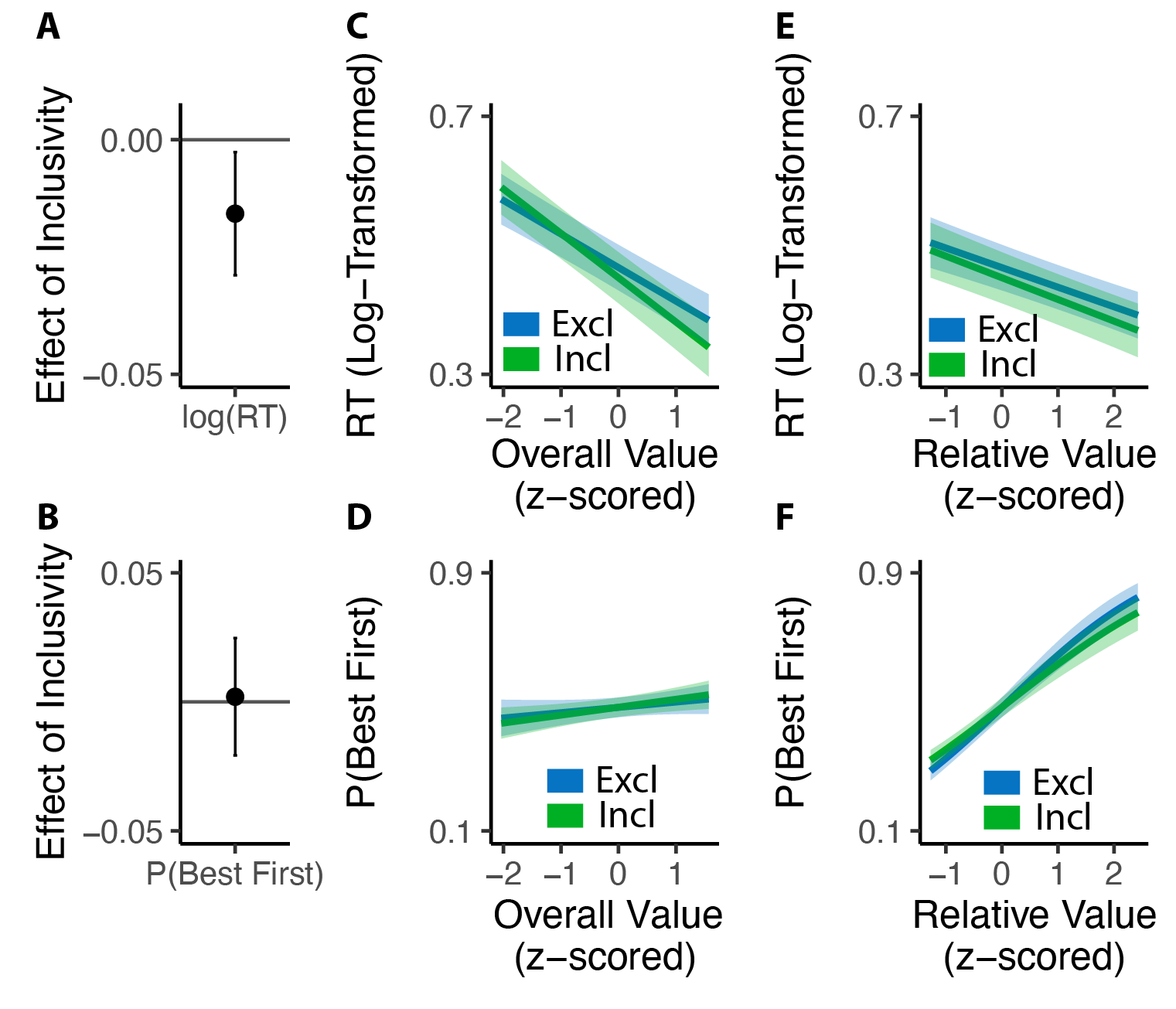

**Figure S5. Influence of inclusivity on choice behavior in Study 3A. (A-F)** The effect of inclusivity on choice behavior is consistent with the behavioral patterns in Study 1 (**Figure 2**).

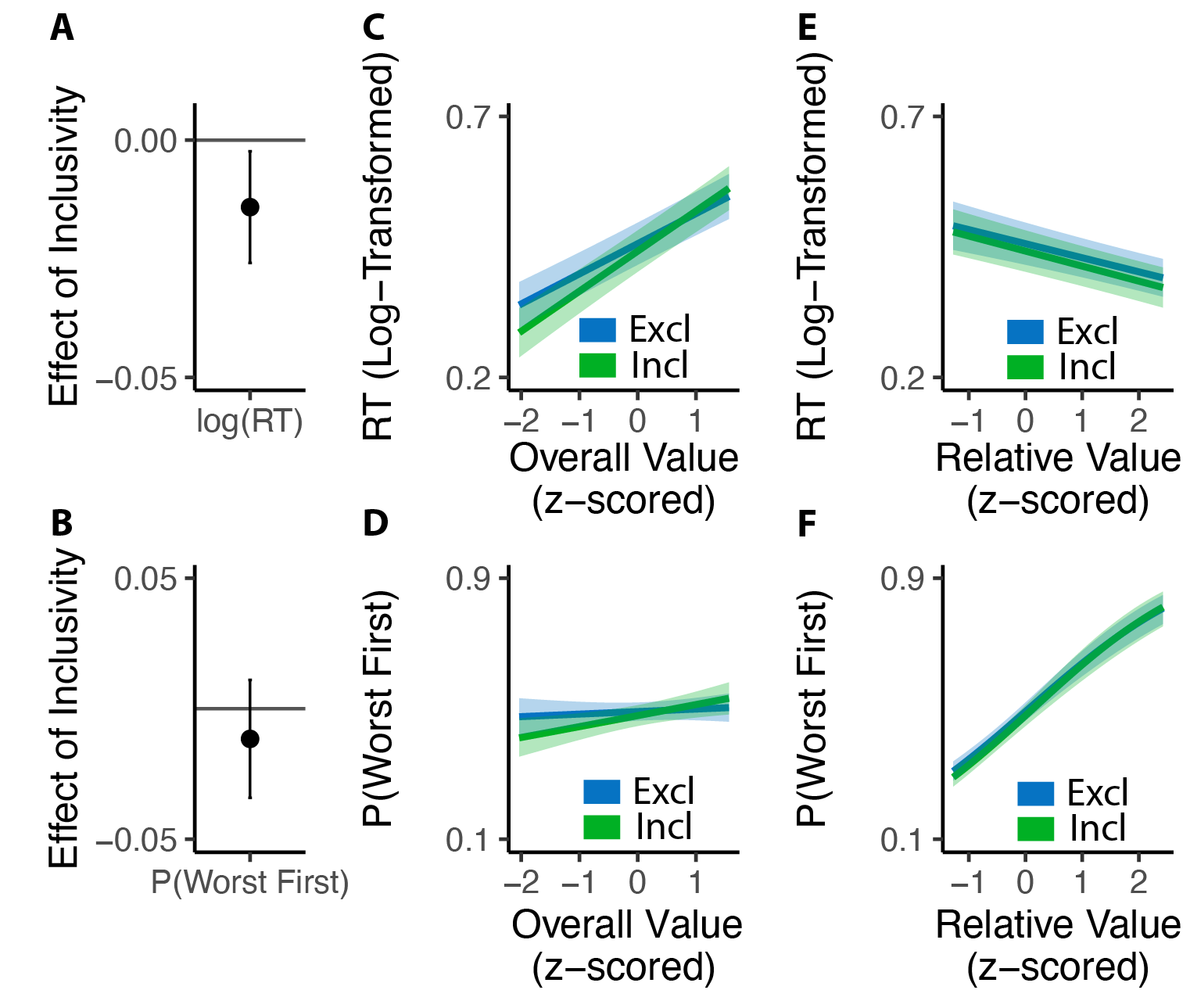

**Figure S6. Influence of inclusivity on choice behavior in Study 3B. (A-F)** The effect of inclusivity on choice behavior is consistent with the behavioral patterns in Study 2 (**Figure 5**).

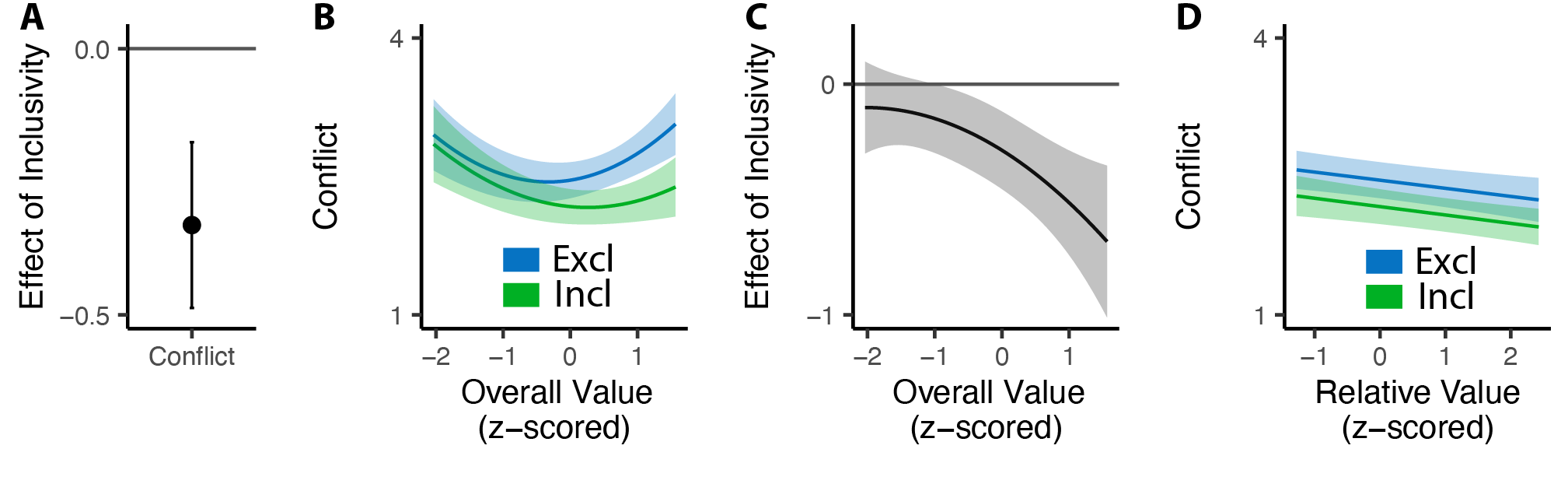
**Figure S7. Influence of inclusivity on choice conflict in Study 3A. (A-D)**The effect of inclusivity on choice conflict replicates the results in Study 1 (**Figure 4**).

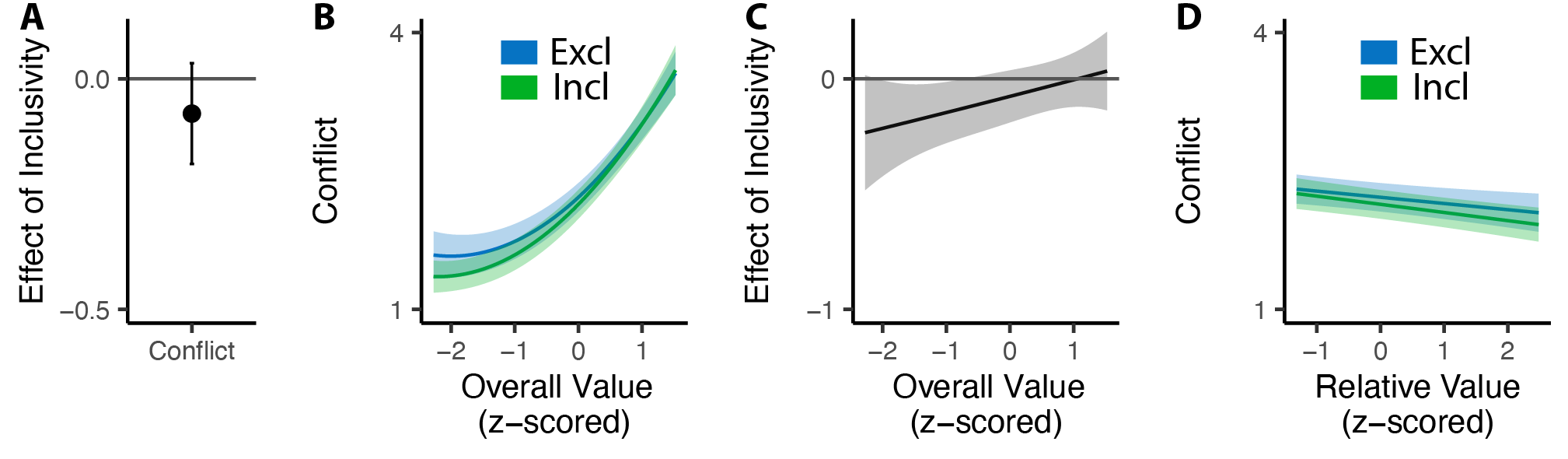

**Figure S8. Influence of inclusivity on choice conflict in Study 3B. (A-D)**The effect of inclusivity on choice conflict replicates the results in Study 2 (**Figure 6**).

### 11. Supplementary figures for Study 4

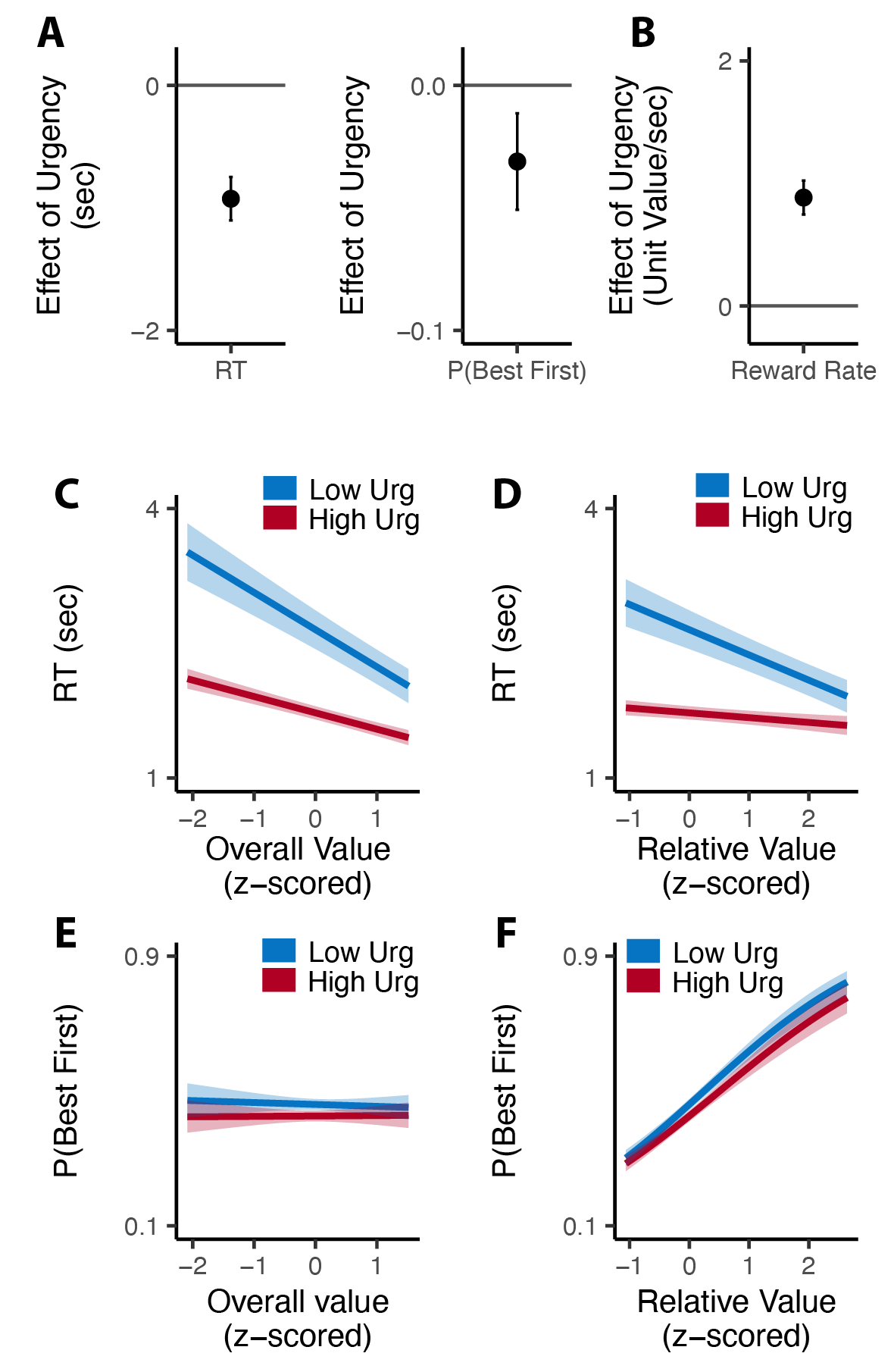

**Figure S9. Influence of urgency on choice behavior in Study 4. (A-B)** People made faster and less accurate choices with high choice urgency, leading to higher reward rate. **(C-D)** Choice urgency similarly modulates the influence of overall value and relative value on choice reaction time. **(E-F)** The effects of overall value and relative value on choice accuracy do not differ across high and low choice urgency.

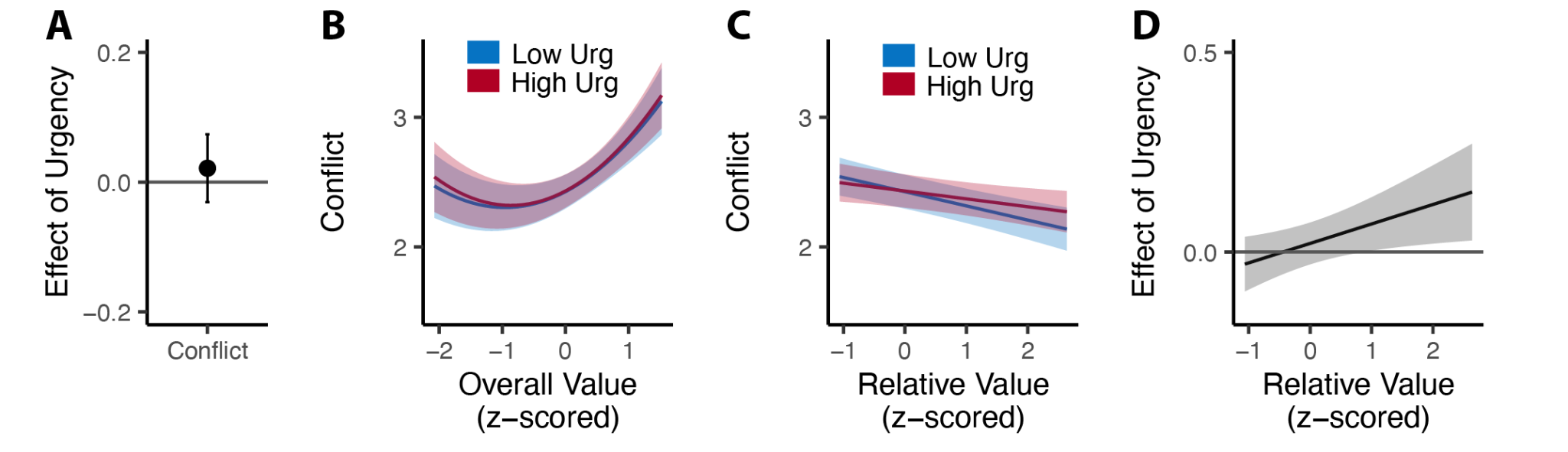

**Figure S10. Influence of urgency on choice conflict in Study 4. (A)** High choice urgency does not influence overall choice conflict. **(B)** Choice conflict increases when the overall value is extremely high or low, regardless of choice urgency. **(C)** With high choice urgency, higher relative value does not reduce choice conflict. **(D)** Higher choice urgency increases choice conflict when the relative value is larger.

### 12. Anova tables (Type III) for mixed-effect models

**Table S14. Anova table for mixed-effect model of RT in Study 1**

|  | Chisq | Df | Pr(>Chisq) |
| --- | --- | --- | --- |
| (Intercept) | 1104.124 | 1 | **0.000** |
| Inclusivity | 53.594 | 1 | **0.000** |
| Overall value | 225.653 | 1 | **0.000** |
| Relative value | 143.504 | 1 | **0.000** |
| Trial Order | 30.187 | 1 | **0.000** |
| Incl:OV | 22.984 | 1 | **0.000** |
| Incl:RV | 0.186 | 1 | 0.666 |

**Table S15. Anova table for mixed-effect model of Accuracy in Study 1**

|  | Chisq | Df | Pr(>Chisq) |
| --- | --- | --- | --- |
| (Intercept) | 3.014 | 1 | 0.083 |
| Inclusivity | 4.779 | 1 | **0.029** |
| Overall value | 0.070 | 1 | 0.791 |
| Relative value | 281.512 | 1 | **0.000** |
| Trial Order | 1.935 | 1 | 0.164 |
| Incl:OV | 2.092 | 1 | 0.148 |
| Incl:RV | 1.348 | 1 | 0.246 |

**Table S16. Anova table for mixed-effect model of conflict in Study 1**

|  | Chisq | Df | Pr(>Chisq) |
| --- | --- | --- | --- |
| (Intercept) | 955.252 | 1 | **0.000** |
| Inclusivity | 32.932 | 1 | **0.000** |
| poly(Overall value, 2) | 15.932 | 2 | **0.000** |
| Relative value | 17.299 | 1 | **0.000** |
| Trial Order | 1.089 | 1 | 0.297 |
| Incl:poly(OV,2) | 16.276 | 2 | **0.000** |
| Incl:RV | 1.369 | 1 | 0.242 |

**Table S17. Anova table for mixed-effect model of RT in Study 2**

|  | Chisq | Df | Pr(>Chisq) |
| --- | --- | --- | --- |
| (Intercept) | 1729.324 | 1 | **0.000** |
| Inclusivity | 17.333 | 1 | **0.000** |
| Overall value | 348.369 | 1 | **0.000** |
| Relative value | 117.882 | 1 | **0.000** |
| Trial Order | 58.055 | 1 | **0.000** |
| Incl:OV | 8.540 | 1 | **0.003** |
| Incl:RV | 0.084 | 1 | 0.772 |

**Table S18. Anova table for mixed-effect model of accuracy in Study 2**

|  | Chisq | Df | Pr(>Chisq) |
| --- | --- | --- | --- |
| (Intercept) | 28.740 | 1 | **0.000** |
| Inclusivity | 11.225 | 1 | **0.001** |
| Overall value | 11.074 | 1 | **0.001** |
| Relative value | 206.063 | 1 | **0.000** |
| Trial Order | 5.170 | 1 | **0.023** |
| Incl:OV | 1.930 | 1 | 0.165 |
| Incl:RV | 3.273 | 1 | 0.070 |

**Table S19. Anova table for mixed-effect model of conflict in Study 2**

|  | Chisq | Df | Pr(>Chisq) |
| --- | --- | --- | --- |
| (Intercept) | 2264.265 | 1 | **0.000** |
| Inclusivity | 4.711 | 1 | **0.030** |
| poly(Overall value, 2) | 137.778 | 2 | **0.000** |
| Relative value | 16.876 | 1 | **0.000** |
| Trial Order | 0.004 | 1 | 0.951 |
| Incl:poly(OV,2) | 6.165 | 2 | **0.046** |
| Incl:RV | 1.669 | 1 | 0.196 |

**Table S20. Anova table for mixed-effect model of log-transformedRT in Study 3A**

|  | Chisq | Df | Pr(>Chisq) |
| --- | --- | --- | --- |
| (Intercept) | 581.571 | 1 | **0.000** |
| Inclusivity | 6.098 | 1 | **0.014** |
| Overall value | 175.201 | 1 | **0.000** |
| Relative value | 84.647 | 1 | **0.000** |
| Trial Order | 48.524 | 1 | **0.000** |
| Incl:OV | 10.459 | 1 | **0.001** |
| Incl:RV | 0.696 | 1 | 0.404 |

**Table S18. Anova table for mixed-effect model of accuracy in Study 3A**

|  | Chisq | Df | Pr(>Chisq) |
| --- | --- | --- | --- |
| (Intercept) | 1.446 | 1 | 0.229 |
| Inclusivity | 0.001 | 1 | 0.972 |
| Overall value | 5.854 | 1 | **0.016** |
| Relative value | 203.222 | 1 | **0.000** |
| Trial Order | 1.783 | 1 | 0.182 |
| Incl:OV | 0.295 | 1 | 0.587 |
| Incl:RV | 3.349 | 1 | 0.067 |

**Table S19. Anova table for mixed-effect model of conflict in Study 3A**

|  | Chisq | Df | Pr(>Chisq) |
| --- | --- | --- | --- |
| (Intercept) | 1064.430 | 1 | **0.000** |
| Inclusivity | 17.363 | 1 | **0.000** |
| poly(Overall value, 2) | 25.416 | 2 | **0.000** |
| Relative value | 16.431 | 1 | **0.000** |
| Trial Order | 0.002 | 1 | 0.966 |
| Incl:poly(OV,2) | 10.745 | 2 | **0.005** |
| Incl:RV | 0.008 | 1 | 0.930 |

**Table S20. Anova table for mixed-effect model of log-transformed RT in Study 3B**

|  | Chisq | Df | Pr(>Chisq) |
| --- | --- | --- | --- |
| (Intercept) | 496.766 | 1 | **0.000** |
| Inclusivity | 6.477 | 1 | **0.011** |
| Overall value | 241.762 | 1 | **0.000** |
| Relative value | 74.952 | 1 | **0.000** |
| Trial Order | 56.848 | 1 | **0.000** |
| Incl:OV | 14.205 | 1 | **0.000** |
| Incl:RV | 0.115 | 1 | 0.734 |

**Table S21. Anova table for mixed-effect model of accuracy in Study 3B**

|  | Chisq | Df | Pr(>Chisq) |
| --- | --- | --- | --- |
| (Intercept) | 1.227 | 1 | 0.268 |
| Inclusivity | 0.915 | 1 | 0.339 |
| Overall value | 4.546 | 1 | **0.033** |
| Relative value | 224.965 | 1 | **0.000** |
| Trial Order | 0.409 | 1 | 0.523 |
| Incl:OV | 3.179 | 1 | 0.075 |
| Incl:RV | 0.294 | 1 | 0.588 |

**Table S22. Anova table for mixed-effect model of conflict in Study 3B**

|  | Chisq | Df | Pr(>Chisq) |
| --- | --- | --- | --- |
| (Intercept) | 1282.169 | 1 | **0.000** |
| Inclusivity | 1.829 | 1 | 0.176 |
| poly(Overall value, 2) | 151.088 | 2 | **0.000** |
| Relative value | 18.064 | 1 | **0.000** |
| Trial Order | 0.003 | 1 | 0.955 |
| Incl:poly(OV,2) | 6.254 | 2 | **0.044** |
| Incl:RV | 0.696 | 1 | 0.404 |

**Table S23. Anova table for mixed-effect model of log-transformed RT in Study 4**

|  | Chisq | Df | Pr(>Chisq) |
| --- | --- | --- | --- |
| (Intercept) | 1023.952 | 1 | **0** |
| Inclusivity | 105.234 | 1 | **0** |
| Overall value | 169.022 | 1 | **0** |
| Relative value | 77.394 | 1 | **0** |
| Trial Order | 49.780 | 1 | **0** |
| Incl:OV | 39.3910 | 1 | **0** |
| Incl:RV | 40.074 | 1 | **0** |

**Table S24. Anova table for mixed-effect model of accuracy in Study 4**

|  | Chisq | Df | Pr(>Chisq) |
| --- | --- | --- | --- |
| (Intercept) | 94.493 | 1 | **0.000** |
| Inclusivity | 9.118 | 1 | **0.003** |
| Overall value | 0.058 | 1 | 0.809 |
| Relative value | 308.287 | 1 | **0.000** |
| Trial Order | 1.121 | 1 | 0.290 |
| Incl:OV | 0.279 | 1 | 0.597 |
| Incl:RV | 0.984 | 1 | 0.321 |

**Table S25. Anova table for mixed-effect model of conflict in Study 4**

|  | Chisq | Df | Pr(>Chisq) |
| --- | --- | --- | --- |
| (Intercept) | 1878.058 | 1 | **0.000** |
| Inclusivity | 0.646 | 1 | 0.422 |
| poly(Overall value, 2) | 47.014 | 2 | **0.000** |
| Relative value | 15.978 | 1 | **0.000** |
| Trial Order | 1.420 | 1 | 0.233 |
| Incl:poly(OV,2) | 0.829 | 2 | 0.661 |
| Incl:RV | 5.343 | 1 | **0.021** |

**Table S26. Anova table for mixed-effect model of log-transformed RT in Study S1**

|  | Chisq | Df | Pr(>Chisq) |
| --- | --- | --- | --- |
| (Intercept) | 946.175 | 1 | **0.000** |
| Inclusivity | 18.721 | 1 | **0.000** |
| Overall value | 109.950 | 1 | **0.000** |
| Relative value | 304.579 | 1 | **0.000** |
| Trial Order | 169.760 | 1 | **0.000** |
| Incl:OV | 11.448 | 1 | **0.001** |
| Incl:RV | 0.787 | 1 | 0.375 |

**Table S27. Anova table for mixed-effect model of accuracy in Study S1**

|  | Chisq | Df | Pr(>Chisq) |
| --- | --- | --- | --- |
| (Intercept) | 38.324 | 1 | **0.000** |
| Inclusivity | 1.448 | 1 | 0.229 |
| Overall value | 3.390 | 1 | **0.066** |
| Relative value | 463.365 | 1 | **0.000** |
| Trial Order | 1.608 | 1 | 0.205 |
| Incl:OV | 0.378 | 1 | 0.539 |
| Incl:RV | 1.981 | 1 | 0.159 |

**Table S28. Anova table for mixed-effect model of conflict in Study S1**

|  | Chisq | Df | Pr(>Chisq) |
| --- | --- | --- | --- |
| (Intercept) | 1222.159 | 1 | **0.000** |
| Inclusivity | 13.203 | 1 | **0.000** |
| poly(Overall value, 2) | 64.673 | 2 | **0.000** |
| Relative value | 15.592 | 1 | **0.000** |
| Trial Order | 0.303 | 1 | 0.582 |
| Incl:poly(OV,2) | 11.435 | 2 | **0.003** |
| Incl:RV | 0.507 | 1 | 0.477 |

**Table S29. Anova table for mixed-effect model of log-transformed RT in the incentivized subset of Study S1**

|  | Chisq | Df | Pr(>Chisq) |
| --- | --- | --- | --- |
| (Intercept) | 529.186 | 1 | **0.000** |
| Inclusivity | 6.035 | 1 | **0.014** |
| Overall value | 30.262 | 1 | **0.000** |
| Relative value | 155.779 | 1 | **0.000** |
| Trial Order | 107.164 | 1 | **0.000** |
| Incl:OV | 4.942 | 1 | **0.026** |
| Incl:RV | 1.352 | 1 | 0.245 |

**Table S30. Anova table for mixed-effect model of accuracy in the incentivized subset of Study S1**

|  | Chisq | Df | Pr(>Chisq) |
| --- | --- | --- | --- |
| (Intercept) | 19.276 | 1 | **0.000** |
| Inclusivity | 1.692 | 1 | 0.193 |
| Overall value | 1.676 | 1 | 0.195 |
| Relative value | 267.716 | 1 | **0.000** |
| Trial Order | 2.022 | 1 | 0.155 |
| Incl:OV | 0.026 | 1 | 0.873 |
| Incl:RV | 0.360 | 1 | 0.549 |

**Table S31. Anova table for mixed-effect model of conflict in the incentivized subset of Study S1**

|  | Chisq | Df | Pr(>Chisq) |
| --- | --- | --- | --- |
| (Intercept) | 697.905 | 1 | **0.000** |
| Inclusivity | 5.985 | 1 | **0.014** |
| poly(Overall value, 2) | 20.565 | 2 | **0.000** |
| Relative value | 9.317 | 1 | **0.002** |
| Trial Order | 0.371 | 1 | 0.542 |
| Incl:poly(OV,2) | 5.716 | 2 | 0.057 |
| Incl:RV | 0.072 | 1 | 0.789 |

**Table S32. Anova table for mixed-effect model of number of chosen options**

**(Study 1 and 3A combined)**

|  | Chisq | Df | Pr(>Chisq) |
| --- | --- | --- | --- |
| (Intercept) | 1767.251 | 1 | **0.000** |
| Study | 0.040 | 1 | 0.841 |
| Conflict | 4.127 | 1 | **0.042** |
| poly(Overall Value, 2) | 391.884 | 2 | **0.000** |
| Trial Order | 2.307 | 1 | 0.129 |
| Relative Value | 20.305 | 1 | **0.000** |
| Initial RT | 60.381 | 1 | **0.000** |
| Initial Accuracy | 17.899 | 1 | **0.000** |
| Study:Conflict | 0.075 | 1 | 0.785 |
| Study:poly(Overall Value, 2) | 3.372 | 2 | 0.185 |
| Study:Trial Order | 0.686 | 1 | 0.408 |
| Study:Relative Value | 0.000 | 1 | 0.987 |
| Study:Initial RT | 32.771 | 1 | **0.000** |
| Study:Initial Accuracy | 0.676 | 1 | 0.411 |

**Table S33. Anova table for mixed-effect model of number of chosen options**

**(Study 2 and 3B combined)**

|  | Chisq | Df | Pr(>Chisq) |
| --- | --- | --- | --- |
| (Intercept) | 3103.744 | 1 | **0.000** |
| Study | 3.187 | 1 | 0.074 |
| Conflict | 30.494 | 1 | **0.000** |
| poly(Overall Value, 2) | 402.373 | 2 | **0.000** |
| Trial Order | 0.583 | 1 | 0.445 |
| Relative Value | 5.916 | 1 | **0.015** |
| Initial RT | 182.820 | 1 | **0.000** |
| Initial Accuracy | 43.452 | 1 | **0.000** |
| Study:Conflict | 1.482 | 1 | 0.224 |
| Study:poly(Overall Value, 2) | 2.637 | 2 | 0.268 |
| Study:Trial Order | 1.187 | 1 | 0.276 |
| Study:Relative Value | 0.032 | 1 | 0.858 |
| Study:Initial RT | 71.589 | 1 | **0.000** |
| Study:Initial Accuracy | 0.044 | 1 | 0.834 |

**Table S34. Anova table for mixed-effect model of ratio of kept suboptimal options**

**(Study 1 and 3A combined)**

|  | Chisq | Df | Pr(>Chisq) |
| --- | --- | --- | --- |
| (Intercept) | 2895.356 | 1 | **0.000** |
| Study | 2.685 | 1 | 0.101 |
| Conflict | 7.254 | 1 | **0.007** |
| N of suboptimal options | 4272.804 | 1 | **0.000** |
| poly(Overall Value, 2) | 22.320 | 2 | **0.000** |
| Trial Order | 4.705 | 1 | **0.030** |
| Relative Value | 274.547 | 1 | **0.000** |
| Initial RT | 2.465 | 1 | 0.116 |
| Initial Accuracy | 109.546 | 1 | **0.000** |
| Study:Conflict | 1.391 | 1 | 0.238 |
| Study:Nsuboptimal | 0.004 | 1 | 0.947 |
| Study:poly(OV, 2) | 5.530 | 2 | 0.063 |
| Study:Trial Order | 4.863 | 1 | **0.027** |
| Study:RV | 6.616 | 1 | **0.010** |
| Study:Initial RT | 1.321 | 1 | 0.250 |
| Study:Initial Accuracy | 0.605 | 1 | 0.437 |

**Table S35. Anova table for mixed-effect model of ratio of kept suboptimal options**

**(Study 2 and 3B combined)**

|  | Chisq | Df | Pr(>Chisq) |
| --- | --- | --- | --- |
| (Intercept) | 1570.737 | 1 | **0.000** |
| Study | 0.201 | 1 | 0.654 |
| Conflict | 10.240 | 1 | **0.001** |
| N of suboptimal options | 3292.247 | 1 | **0.000** |
| poly(Overall Value, 2) | 96.996 | 2 | **0.000** |
| Trial Order | 1.835 | 1 | 0.175 |
| Relative Value | 261.614 | 1 | **0.000** |
| Initial RT | 26.380 | 1 | **0.000** |
| Initial Accuracy | 95.980 | 1 | **0.000** |
| Study:Conflict | 1.343 | 1 | 0.247 |
| Study:Nsuboptimal | 0.721 | 1 | 0.396 |
| Study:poly(OV, 2) | 0.156 | 2 | 0.925 |
| Study:Trial Order | 2.756 | 1 | 0.097 |
| Study:RV | 2.339 | 1 | 0.126 |
| Study:Initial RT | 17.895 | 1 | **0.000** |
| Study:Initial Accuracy | 0.226 | 1 | 0.635 |

**Table S36. Anova table for mixed-effect model of number of kept options**

**(Study 1, 2 and 3 combined)**

|  | Chisq | Df | Pr(>Chisq) |
| --- | --- | --- | --- |
| (Intercept) | 4862.456 | 1 | **0.000** |
| Study (1&2 vs 3) | 2.078 | 1 | 0.149 |
| Action (selection - removal) | 14.591 | 1 | **0.000** |
| Conflict | 30.308 | 1 | **0.000** |
| poly(Overall Value, 2) | 733.327 | 2 | **0.000** |
| Trial Order | 3.285 | 1 | 0.070 |
| Relative Value | 5.644 | 1 | **0.018** |
| Initial RT | 0.310 | 1 | 0.578 |
| Initial Accuracy | 0.635 | 1 | 0.426 |
| Study:Action | 0.007 | 1 | 0.931 |
| Study:Conflict | 0.955 | 1 | 0.328 |
| Study:poly(OV, 2) | 5.409 | 2 | 0.067 |
| Study:Trial Order | 1.917 | 1 | 0.166 |
| Study:RV | 0.010 | 1 | 0.922 |
| Study:Initial RT | 0.851 | 1 | 0.356 |
| Study:Initial Accuracy | 0.665 | 1 | 0.415 |
| Action:Conflict | 4.711 | 1 | **0.030** |
| Action:poly(OV, 2) | 66.075 | 2 | **0.000** |
| Action:Trial Order | 2.951 | 1 | 0.086 |
| Action:RV | 46.028 | 1 | **0.000** |
| Action:Initial RT | 200.229 | 1 | **0.000** |
| Action:Initial Accuracy | 59.773 | 1 | **0.000** |
| Study:Action:Conflict | 0.205 | 1 | 0.651 |
| Study:Action:poly(OV, 2) | 1.302 | 2 | 0.521 |
| Study:Action:Trial Order | 0.770 | 1 | 0.380 |
| Study:Action:RV | 0.803 | 1 | 0.370 |
| Study:Action:Initial RT | 106.536 | 1 | **0.000** |
| Study:Action:Initial Accuracy | 0.043 | 1 | 0.836 |

**Table S37. Anova table for mixed-effect model of ratio of kept suboptimal options**

**(Study 1, 2 and 3 combined)**

|  | Chisq | Df | Pr(>Chisq) |
| --- | --- | --- | --- |
| (Intercept) | 3627.441 | 1 | **0.000** |
| Study (1&2 vs 3) | 0.055 | 1 | 0.815 |
| Action (selection - removal) | 55.526 | 1 | **0.000** |
| Conflict | 19.043 | 1 | **0.000** |
| N of suboptimal options | 5156.575 | 1 | **0.000** |
| poly(Overall Value, 2) | 50.807 | 2 | **0.000** |
| Trial Order | 6.935 | 1 | **0.008** |
| Relative Value | 535.953 | 1 | **0.000** |
| Initial RT | 24.277 | 1 | **0.000** |
| Initial Accuracy | 204.646 | 1 | **0.000** |
| Study:Action | 0.122 | 1 | 0.727 |
| Study:Conflict | 0.084 | 1 | 0.772 |
| Study:Nsuboptimal | 2.131 | 1 | 0.144 |
| Study:poly(OV, 2) | 0.893 | 2 | 0.640 |
| Study:Trial Order | 6.485 | 1 | **0.011** |
| Study:RV | 9.562 | 1 | **0.002** |
| Study:Initial RT | 3.633 | 1 | 0.057 |
| Study:Initial Accuracy | 0.469 | 1 | 0.493 |
| Action:Conflict | 1.195 | 1 | 0.274 |
| Action:Nsuboptimal | 2.584 | 1 | 0.108 |
| Action:poly(OV, 2) | 138.196 | 2 | **0.000** |
| Action:Trial Order | 0.206 | 1 | 0.650 |
| Action:RV | 11.998 | 1 | **0.001** |
| Action:Initial RT | 6.403 | 1 | **0.011** |
| Action:Initial Accuracy | 11.215 | 1 | **0.001** |
| Study:Action:Conflict | 2.508 | 1 | 0.113 |
| Study:Action:Nsuboptimal | 0.609 | 1 | 0.435 |
| Study:Action:poly(OV, 2) | 0.298 | 2 | 0.862 |
| Study:Action:Trial Order | 0.004 | 1 | 0.948 |
| Study:Action:RV | 3.282 | 1 | 0.070 |
| Study:Action:Initial RT | 13.740 | 1 | **0.000** |
| Study:Action:Initial Accuracy | 0.008 | 1 | 0.927 |
